# Spatio-temporal Control of ERK Pulse Frequency Coordinates Fate Decisions during Mammary Acinar Morphogenesis

**DOI:** 10.1101/2020.11.20.387167

**Authors:** Pascal Ender, Paolo Armando Gagliardi, Maciej Dobrzyński, Agne Frismantiene, Coralie Dessauges, Thomas Höhener, Marc-Antoine Jacques, Andrew R. Cohen, Olivier Pertz

**Affiliations:** Institute of Cell Biology, University of Bern, Baltzerstrasse 4, 3012 Bern, Switzerland; Department of Electrical and Computer Engineering, Drexel University, 3120-40 Market Street, Suite 313, Philadelphia, PA 19104, USA

## Abstract

The signaling events controlling proliferation, survival, and apoptosis during mammary epithelial acinar morphogenesis remain poorly characterized. By imaging single-cell ERK activity dynamics in MCF10A acini, we find that these fates depend on the average frequency of non-periodic ERK pulses. High pulse frequency is observed during initial acinus growth, correlating with rapid cell motility. Subsequent decrease in motility correlates with lower ERK pulse frequency and quiescence. Later, during lumen formation, coordinated ERK waves emerge across multiple cells of an acinus, correlating with high and low ERK pulse frequency in outer surviving and inner dying cells respectively. Optogenetic entrainment of ERK pulses causally connects high ERK pulse frequency with inner cell survival. Acini harboring the PIK3CA H1047R mutation, commonly observed in breast cancer, display increased ERK pulse frequency, inner cell survival and loss of lumen formation. Thus, fate decisions during acinar morphogenesis are fine-tuned by different spatio-temporal coordination modalities of ERK pulse frequency.

## Introduction

Mammary organogenesis involves formation of a rudimentary gland during embryogenesis, followed by proliferation and branching invasion led by multi-layered terminal end buds (TEBs) during puberty. Cells in the inner TEB layers then undergo apoptosis to form the ductal lumen. During pregnancy, secretory alveoli are then formed at the ends of the ductal tree (Inman et al., 2015; Paine and Lewis, 2017). Morphogenesis of this complex structure requires spatial and temporal control of cell fates such as proliferation, survival, migration and death. However, the spatio-temporal signaling events that regulate such fate decisions remain poorly explored. The epidermal growth factor (EGF) receptor (EGFR) – mitogen activated protein kinase (MAPK) signaling cascade, that ultimately leads to activation of the extracellular regulated kinase (ERK) is a key pathway involved in mammary gland development. EGFR-ERK signaling results in the upregulation of gene products involved in a wide variety of processes such as proliferation, survival, migration and differentiation (Lavoie et al., 2020). EGFR activity is required for mammary gland morphogenesis in mice (Sebastian et al., 1998). Paracrine amphiregulin release by the matrix metallo-protease (MMP) ADAM-17 and its binding to EGFR mediates the effects of estrogen receptor α to promote mammary gland development and growth (Ciarloni et al., 2007; Sternlicht et al., 2005). EGFR-dependent ERK activity is enriched at the front of elongating tubes and coordinates cell migration (Huebner et al., 2016). In 3D mammary acini, oncogenic ERK activation suppresses apoptosis and thus lumen formation (Reginato et al., 2005). In primary mammary cell culture, ERK activity is also crucial for survival (Finlay et al., 2000).

Recent single-cell measurements of ERK activity dynamics in a variety of 2D epithelial monolayer cultures have revealed the existence of non-periodic ERK pulses of constant amplitude and duration (Aikin et al., 2020; Albeck et al., 2013; Aoki et al., 2013; Gagliardi et al., 2021; Goglia et al., 2020; Hino et al., 2020; Hiratsuka et al., 2015). These ERK pulses originate from MAPK network properties such as ultrasensitivity (leading to steep ERK activation at a threshold EGFR input), and negative feedback (leading to ERK adaptation) (Kholodenko et al., 2010; Sparta et al., 2015). An emerging theme is that the average frequency of these non-periodic ERK pulses, referred to as ERK frequency from now on, control apoptosis (low frequency), survival (medium frequency) or proliferation (high frequency) (Albeck et al., 2013; Aoki et al., 2013; Gagliardi et al., 2021). These ERK pulses can either exhibit a stochastic spatially uncorrelated behavior (Albeck et al., 2013; Goglia et al., 2020), or can be organized as wave patterns that regulate collective cell migration in a wound (Aoki et al., 2017; Hino et al., 2020; Hiratsuka et al., 2015), cell survival around sites of apoptotic cell extrusion (Gagliardi et al., 2021; Valon et al., 2021), or extrusion of cancer cells (Aikin et al., 2020). These ERK signaling patterns consist of trigger waves that involve sequential activation of ERK pulses in adjacent epithelial cells through paracrine signaling involving MMP-mediated cleavage of pro-EGF ligands.

In this work, we explore single cell ERK dynamics in 3D mammary acini. Culturing mammary MCF10A cells in a basement membrane matrix (Matrigel) produces acini that retain key features of in vivo breast alveoli. Based on previous work (Anderson et al., 2010; Debnath et al., 2003), we subdivided this process into four stages as depicted in Figure S1A. Stage 1 is characterized by high proliferation rates and basement membrane deposition; stage 2 consists of a quiescent state and presence of an outer cell layer with clear basolateral polarity; stage 3 consists of apoptosis of inner cells that allows formation of a hollow lumen in stage 4. The whole process takes approximately two weeks.

We document different spatio-temporal modalities of ERK signaling during different developmental stages of MCF10A acini formation. Stage 1 is characterized by high ERK frequency, robust proliferation and rapid collective motility. This is followed by a transition to lower ERK frequency and slower collective motility. During stage 2, formation of ERK wave patterns correlates with domains of different ERK frequencies: outer/inner cells display medium/low ERK frequencies, respectively controlling survival and apoptosis. Optogenetic control of ERK signaling shows that ERK pulses control collective migration during stage 1, and that a critical ERK frequency is necessary for survival during stage 2. We characterize a crosstalk between phospho-inositide-3 kinase (PI3K) and MAPK/ERK signaling that feeds into the regulation of ERK frequency. This provides insight into how oncogenic PI3K signaling crosstalks with ERK to contribute to loss of lumen formation. Our work reveals how spatio-temporal control of ERK frequency organizes mammary acinar morphogenesis.

## Results

### Stage 1 proliferative acini exhibit non-periodic MMP/EGFR-dependent ERK pulses whose frequency correlates with collective cell migration speed

To explore single-cell ERK dynamics during acinar morphogenesis, we created MCF10A reporter lines expressing the nuclear marker histone 2B (H2B) fused to miRFP703 with either the ERK activity biosensor ERK-KTR fused to mTurquoise2, or a mCherry-geminin S/G2/M cell cycle marker (Sakaue-Sawano et al., 2017). ERK-KTR reports on ERK activity through reversible nucleus/cytosol shuttling after its phosphorylation by active ERK (Regot et al., 2014) (Figure 1A). These lines were then used to grow acini according to a modified version of a previously described protocol (Debnath et al., 2003). After three days, in which EGF, serum and insulin were required for initial acinar growth, these growth factors (GFs) were removed to study ERK signaling dynamics intrinsic to acinar morphogenesis. Using the geminin marker and a fluorogenic caspase substrate, we evaluated if our protocol recapitulated the proliferation, quiescence and apoptosis fates previously documented during acinar morphogenesis (Figure S1B-E) (Debnath et al., 2002; Liu et al., 2012). Stage 1 acini (4 days post-seeding) displayed elevated levels of proliferation and absence of apoptosis. Stage 2 and 3 acini (7 and 11 days post-seeding) revealed quiescent cells and increased apoptosis leading to lumen formation. Stage 4 acini (14 days post-seeding) exhibited a mature lumen, abundant apoptotic debris, and a small increase in proliferation.

**Figure 1.**
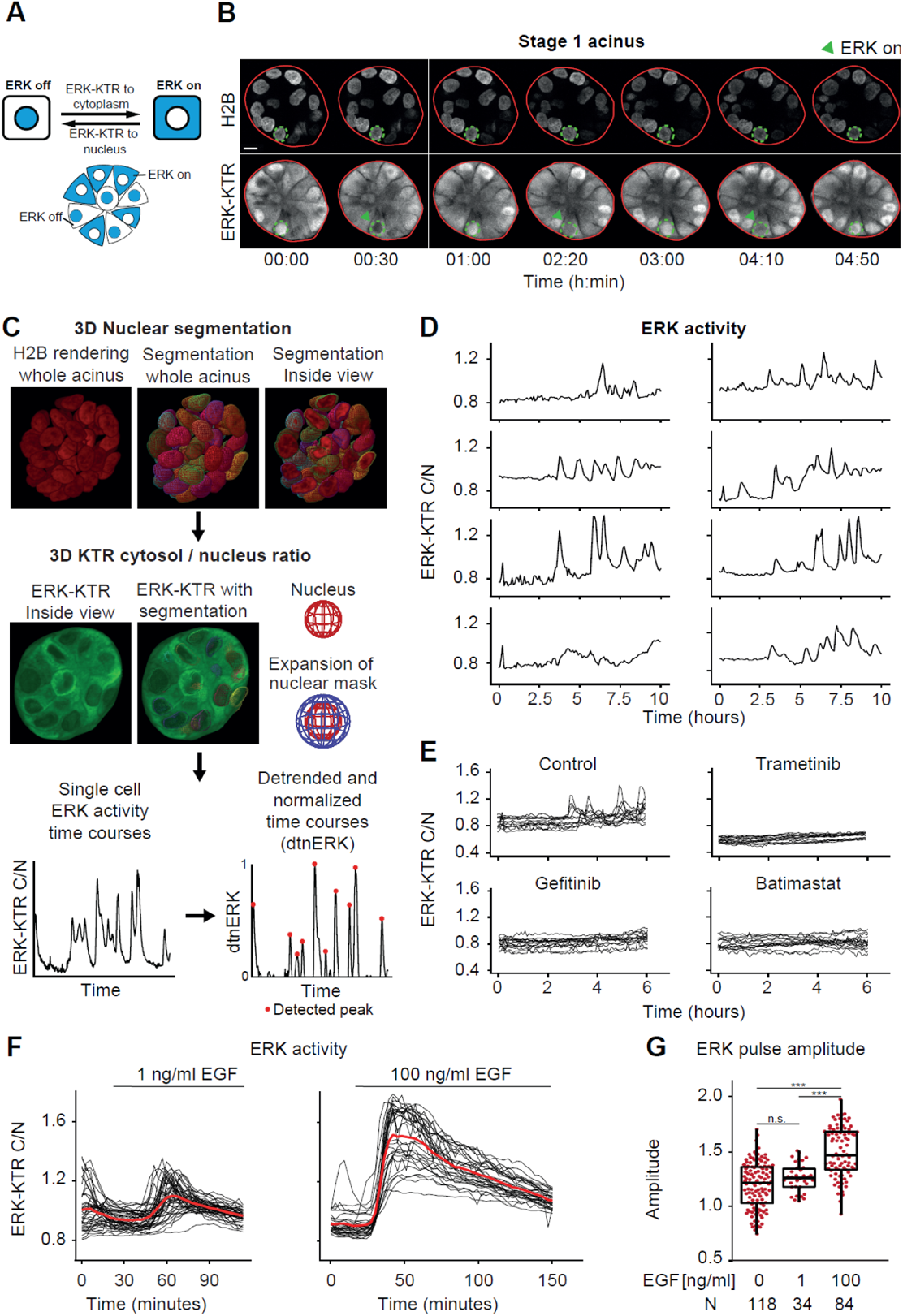
Spontaneous EGFR/MMP-dependent ERK pulses in acini. (A) Schematic representation of the ERK activity - dependent subcellular localization of ERK-KTR. (B) Time-series of the equatorial optical section of an acinus (red plain line) expressing fluorescent H2B and ERK-KTR. Highlighted is a cell (green dotted line) that displays spontaneous ERK activity pulses (arrowheads) resulting in the nuclear to cytoplasmic translocation of ERK-KTR. Scale bar = 10 μm. (C) Image analysis pipeline to extract ERK trajectories from 3D time lapse datasets. Nuclei are segmented and tracked by LEVERJS based on the H2B signal. Single-cell ERK activity levels are the ratio of the median ERK-KTR signal pixel intensities in the voxel mask around the nucleus and the one of the segmented nuclear volume. ERK pulses are detected on detrended ERK trajectories normalized to [0,1]. (D) Representative single-cell ERK trajectories from one acinus. (E) Overlayed ERK trajectories from control and drug-treated acini. Control trajectories correspond to the same acinus as in (D). (F) ERK trajectories from acini treated with 1 or 100 ng/ml EGF at the indicated time points. (G) ERK pulse amplitudes in cells from control and EGF-treated acini. Wilcoxon tests (n.s., P > 0.05; ***, P < 0.001).

We then imaged single-cell ERK dynamics in stage 1 acini using confocal spinning disk microscopy of both the H2B and ERK-KTR channels with time resolutions of 3 - 5 minutes, until acini started to suffer from phototoxicity (observed after 10 - 23 hours). Cells in stage 1 acini displayed asynchronous, non-periodic ERK pulses (Figure 1B) as observed in 2D culture (Aikin et al., 2020; Albeck et al., 2013; Gagliardi et al., 2021). To extract single-cell ERK activity trajectories, we used a customized version of the open-source LEVERJS software (Wait et al., 2014; Winter et al., 2016) that segments and tracks nuclei based on H2B signal; and calculates ERK activity as a ratio of ERK-KTR fluorescence intensities in cytosolic and nuclear voxel masks (Figure 1C). Detrending of the ERK trajectories and normalization of the values to [0,1] generated a reliable input for automated detection of ERK pulses (Figure 1C). Single cell ERK trajectories revealed spontaneous ERK pulses with slightly different amplitudes (Figure 1D). Trametinib-mediated MEK, gefitinib-mediated EGFR, as well as Batimastat-mediated MMP inhibition abolished ERK pulses (Figure 1E). MEK or EGFR inhibition for multiple days led to massive cell death and disintegration of the acini (Figure S2A), suggesting that ERK provides a pro-survival signal. Acute stimulation with 1 ng/ml EGF induced ERK pulses of amplitudes similar to those of spontaneous pulses, while 100 ng/ml EGF induced ERK pulses of higher amplitudes (Figure 1F,G). These results document spontaneous, asynchronous EGFR- and MMP-depedent ERK pulses in stage 1 acini.

As previously described (Wang et al., 2013), stage 1 acini displayed collective cell motility correlating with a rotational movement of 360 degrees of whole acini over multiple hours. Later during stage 1, we observed a transition to a state of slower motility (Figure 2A, Movie S1). This was shown to correlate with deposition of basement membrane around the acinus (Wang et al., 2013). The reduction in migration speed correlated with decreased ERK frequency (Figure 2B-D and S3A, Movie S2), without having a significant effect on ERK pulse amplitude and duration (Figure S3B,C).

**Figure 2.**
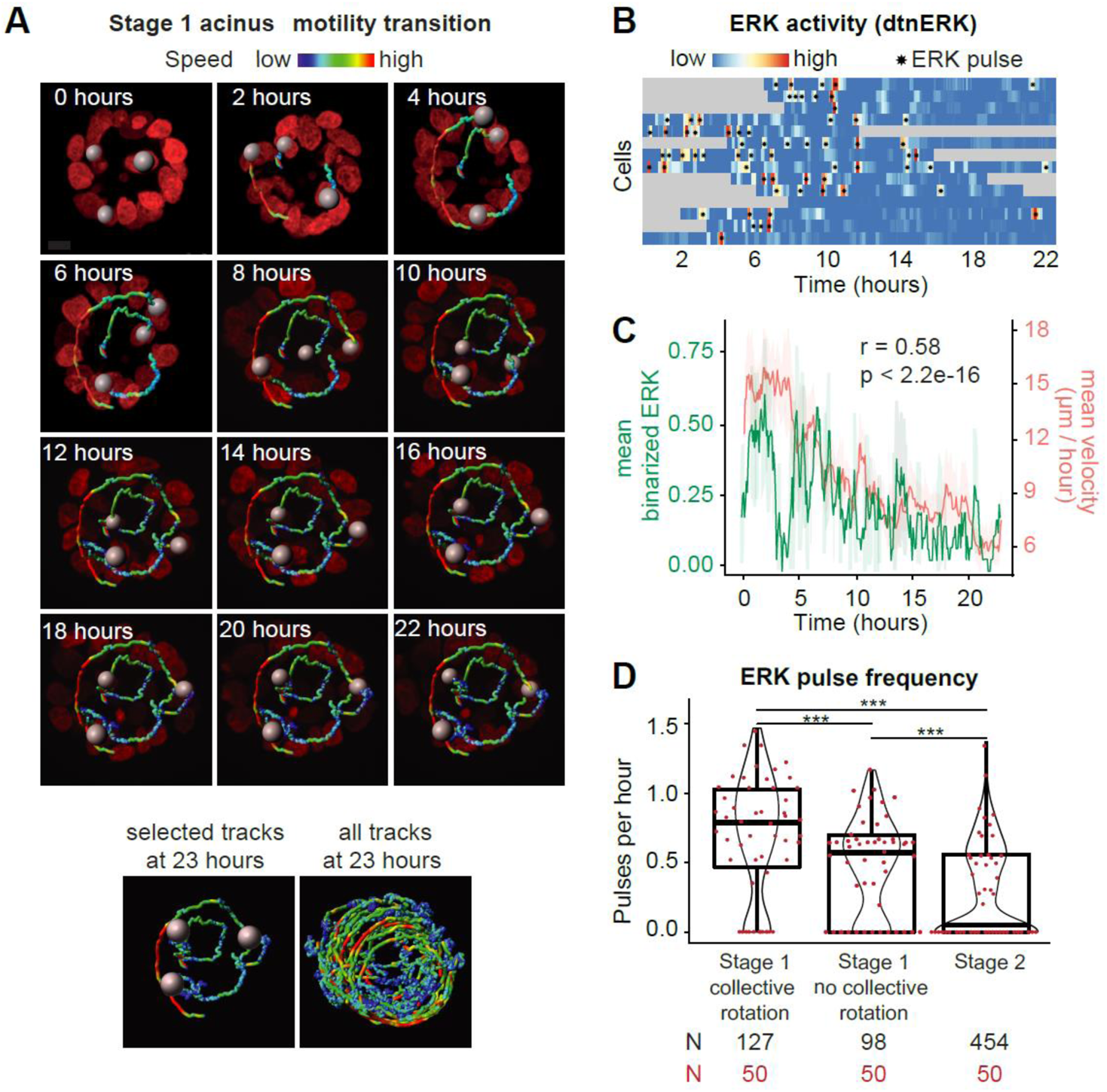
Collective cell migration and ERK pulsing in stage 1 acini. (A) Time series renderings of the cross section of an acinus transitioning from the rapid motility to the slow motility stage. Nuclei and motility tracks color-coded by instantaneous velocity are shown. Scale bar = 10 μm. (B) Analysis of ERK activity in the acinus from (A). Heatmap shows detrended and normalized single cell ERK activity levels over time. Gray areas correspond to time points when a cell was not within the imaged volume. Asterisks indicate individual ERK pulses. (C) Analysis of motility and ERK activity in the acinus from (A). Graph shows mean binarized ERK activity and mean instantaneous velocity with 95% confidence intervals of all imaged cells over time and their Pearson correlation coefficient. Mean binarized ERK activity is used as a measure for the fraction of the cell population in a state of active ERK. (D) ERK pulse frequency from trajectories at different developmental timepoints. Trajectories pooled from 7 (stage 1 rotation), 5 (stage 1 no rotation) and 11 (stage 2) acini. Wilcoxon tests (***, P < 0.001).

### Stage 2 quiescent acini exhibit different ERK frequencies in inner and outer cells, which emerges from collective waves of ERK pulses

We then evaluated ERK dynamics in larger, stage 2 quiescent acini that are characterized by an outer layer of polarized cells, and a less organized inner cell mass destined for apoptosis for future lumen formation (Debnath et al., 2002). When comparing low-motility stage 1 and stage 2 acini, we observed a further reduction in median ERK frequency (Figure 2D), while ERK pulse amplitudes and durations remained almost identical (Figure S3B,C). This change in ERK frequency resulted from a bimodal distribution in which a part of the cell population did not display any ERK pulses (Figure 2D). Because ERK pulse frequency can regulate proliferation, survival and apoptosis fates in MCF10A cells (Aikin et al., 2020; Albeck et al., 2013; Gagliardi et al., 2021), and because inner cells in stage 2 acini are destined to undergo apoptosis, we evaluated ERK pulse frequencies in inner versus outer cells. Outer cells exhibited a significantly higher ERK frequency than inner cells, with the latter often not exhibiting ERK pulses at all (Figure 3A,B). Similar ERK activity amplitude and duration were found in inner/outer cells (Figure S3D,E). Together with our characterization of fate decisions (Figure S1), these results suggest a spatio-temporal mechanism that controls survival versus apoptosis fates through regulation of ERK frequency.

**Figure 3.**
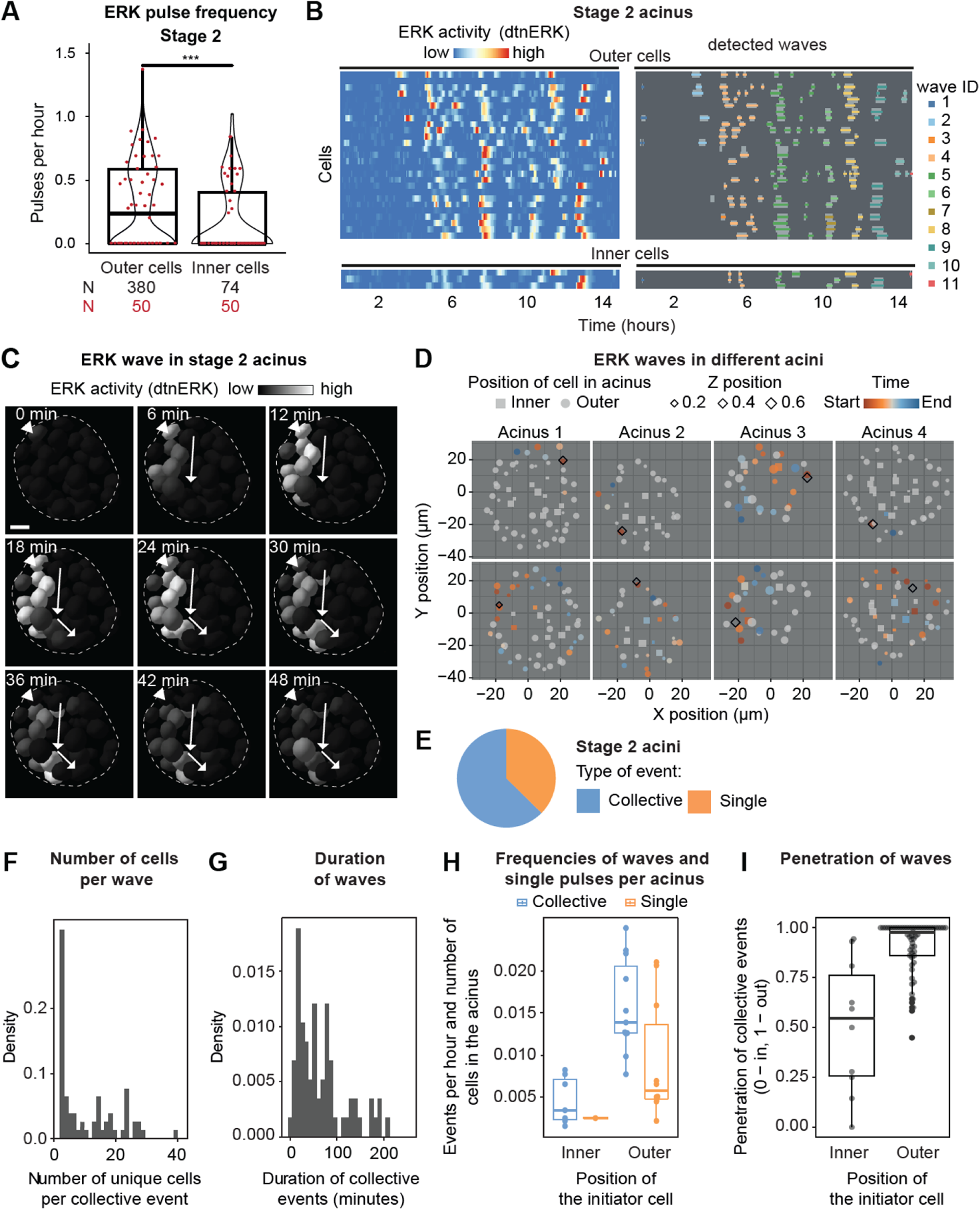
Different ERK pulse frequencies in inner and outer acinar cell layers in stage 2, and collective waves of ERK pulses. (A) ERK pulse frequency from trajectories of cells located in inner versus outer acini layers. Trajectories pooled from 11 acini. Wilcoxon test (***, P < 0.001). (B) Left: heatmap of detrended/normalized single-cell ERK trajectories in outer and inner cells of a representative acinus. Right: detection of individual ERK activity waves in the same acinus. (C) Representative time-series micrographs of ERK wave ID 10 in (B). Nuclei color-coded by ERK-KTR ratios. Arrows depict wave directionality. The arrowhead indicates the initiator cell. Dashed line indicates the acinus border. Scale bar = 10 μm. (D) 2D projection representations of isolated ERK waves from four different acini. Cells that participate in the wave are color-coded by their relative time of activation. Size represents the relative Z position of the cell and shape if they belong to the inner or outer cell population. Top and bottom panels depict two isolated ERK waves for each acinus. (E) Percentage of initiator events that remain restricted to a single cell vs those that lead to collective events. (F) Total numbers of unique cells involved in individual collective events. (G) Durations of individual collective events. (H) Average frequency of single and collective events per acinus, normalized by the number of cells in the acinus. (I) Penetration of collective events across acini. Calculated as the time-averaged fraction of localization of a collective event between the inner (0) and outer (1) cell layer.

A striking feature of stage 2 acini was that they exhibited spatially correlated ERK pulses in the form of waves spreading across multiple cells (Figure 3B,C, Movie S3). We devised computational methods (Figure S3F-H, Materials and Methods) to detect, track, and extract features that describe ERK waves. These ERK waves were observed in all of the stage 2 acini that we imaged (N=11), and exhibited different geometries (Figure 3D). While some ERK pulses remained restricted to single cells, most of the ERK pulses occurred within collective waves (Figure 3E). ERK waves typically involved a median of 6 cells for a median duration of 54 minutes (Figure 3F,G). However, a large variance was observed with some ERK waves involving as little as 2 and as many as 39 cells (almost the whole acinus). ERK waves, as well as isolated ERK pulses, were predominantly initiated in the outer cell layer (Figure 3H). Further, ERK waves that originated in the outer layer displayed a higher bias to remain at that location than those originating in inner cells (Figure 3I). We have previously shown that in 2D MCF10A cultures, as well as in acini, apoptotic cells trigger ERK waves in their neighboring cells (Gagliardi et al., 2021). However, the ERK waves we observed here only rarely coincided with apoptotic events, suggesting that they originate through a different mechanism. Our results strongly suggest that ERK waves contribute to spatially position different ERK pulse frequencies in inner and outer acinar cells.

### Optogenetic control of ERK frequency regulates collective motility, survival and apoptosis fates

To explore the role of ERK pulse frequency during stage 1 collective motility, as well as stage 2 apoptosis and survival fates required for lumen morphogenesis, we used two optogenetic actuators to evoke different ERK pulse frequencies in acini (Figure 4A). Optogenetic fibroblast growth factor receptor (optoFGFR) is a Cry2-based light-activatable receptor tyrosine kinase that activates ERK, Akt and calcium signaling (Kim et al., 2014). OptoRaf is a CIBN/Cry2-based system in which a catalytic Raf domain is recruited to a plasma membrane targeted anchor in a light-dependent fashion to specifically activate ERK (Aoki et al., 2017). We generated stable lines expressing any of the two optogenetic constructs, a spectrally compatible ERK-KTR-mRuby, and H2B-miRFP. Application of a single blue light pulse evoked a discrete optoFGFR- or optoRaf-mediated ERK pulse with similar shape and duration as spontaneous ones (Figure 4B).

**Figure 4.**
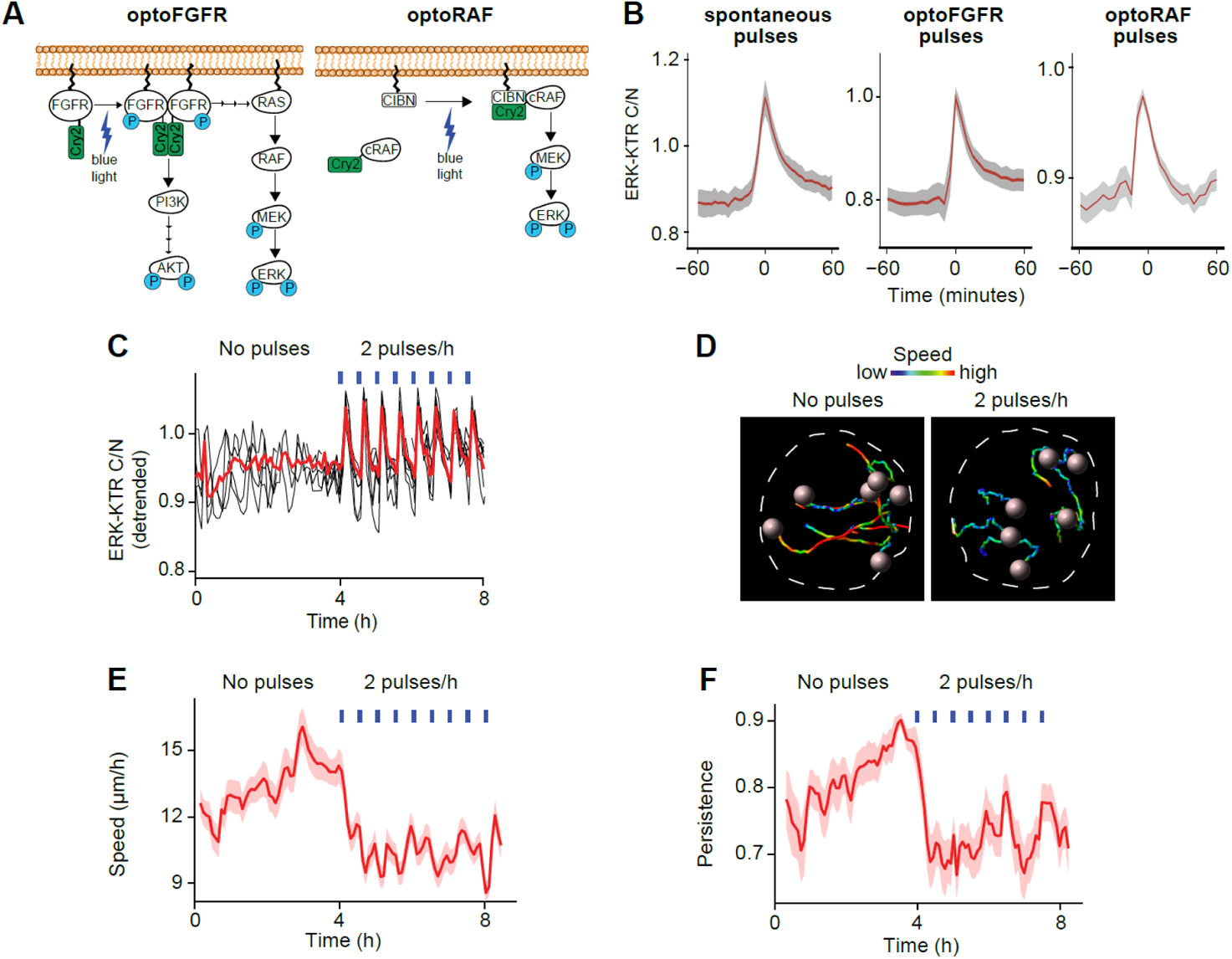
Optogenetic stimulation of acini controls 3D migration properties in stage 1 acini. (A) Cartoon of the optoFGFR and optoRAF systems. OptoFGFR consists of the intracellular domain of FGFR1 linked to the plasma membrane and a Cry2 PHR domain which dimerizes upon blue light stimulation, leading to receptor autophosphorylation and activation of downstream cascades. OptoRAF consists of a Cry2 linked to cRaf and a membrane-linked CIBN domain. CIBN and Cry2 dimerize upon blue light stimulation which recruits cRaf to the plasma membrane where it phosphorylates MEK. (B) Average ERK trajectories from isolated spontaneous and optogenetically induced ERK pulses with 95% confidence interval. Time = 0 corresponds to maximal amplitude of peaks. (C) Overlayed detrended ERK activity trajectories from a stage 1 rotating acinus expressing optoFGFR and stimulated every 30 min starting from 4 hours. Vertical blue lines indicate the blue light stimulation. (D) Six single-cell migration trajectories from the same example organoid color coded according to migration speed. Spheres represent the nuclei positions at the end of the trajectory. The micrographs were taken at 4 and 8 hours of the experiment, each one with the migration trajectories of the past 4 hours. Dashed line indicates the acinar border. (E) Speed and (F) persistence of single-cell migration in the same acinus. Population average and 90% confidence interval are shown.

Since ERK pulse wave patterns can coordinate collective cell migration through regulation of myosin activity (Aoki et al., 2017; Hino et al., 2020), we hypothesized that the asynchronous ERK pulses we observed might coordinate the collective motility pattern in stage 1 acini. We therefore sought to disrupt this process by synchronizing ERK pulses across all the cells of an acinus. We imaged rotating stage 1 acini for 4 hours in the absence of blue light, and observed asynchronous ERK pulses (Figure 4C). We then applied high frequency light pulses at 30 minute intervals, which synchronized ERK pulses across cells (Figure 4C, Movie S4). This switch to synchronous high-frequency ERK pulsing in all cells immediately resulted in decreased cell migration speed and persistence (Figure 4D-F). This suggest that the asynchronous high frequency ERK pulses organize collective cell migration in stage 1 acini.

Next, we tested the hypothesis that the different ERK frequencies observed between outer and inner cells in stage 2 acini regulate survival vs apoptosis cell fates. Using optoFGFR and optoRaf, we evoked frequency-modulated, population-synchronous ERK dynamics in all the cells of an acinus (Figure 5A, Movie S5). Endogenous collective ERK pulses were however still occurring. Because the regulation of apoptosis/survival fates required for lumen formation spans over one week, we could not use our live cell imaging platform to study this process. We therefore used LITOS (LED Illumination Tool for Optogenetic Stimulation) (Höhener et al., 2022) to evoke frequency-modulated ERK pulse regimes in multiwell plates in a tissue culture incubator (Figure S4A). This system evoked similar ERK pulses as observed in our live cell imaging system (Figure S4B). We then stimulated stage 2 acini with light pulses delivered at different frequencies for 7 days and scored the distribution of acini that exhibited filled lumen, partially cleared lumen or cleared lumen (Figure 5B). Using both optoFGFR and optoRaf, we observed that ERK pulses induced every 0.5, 1, 2, 3, 4 but not 10 hours led to survival of inner cells, increasing the number of acini with filled or partially cleared lumina (Figure 5C). These results further suggest that survival and apoptosis fates are regulated by a frequency encoded ERK signal. The optoRaf experiments indicate that high frequency ERK pulsing alone is sufficient to induce survival independently of PI3K/Akt signaling.

**Figure 5.**
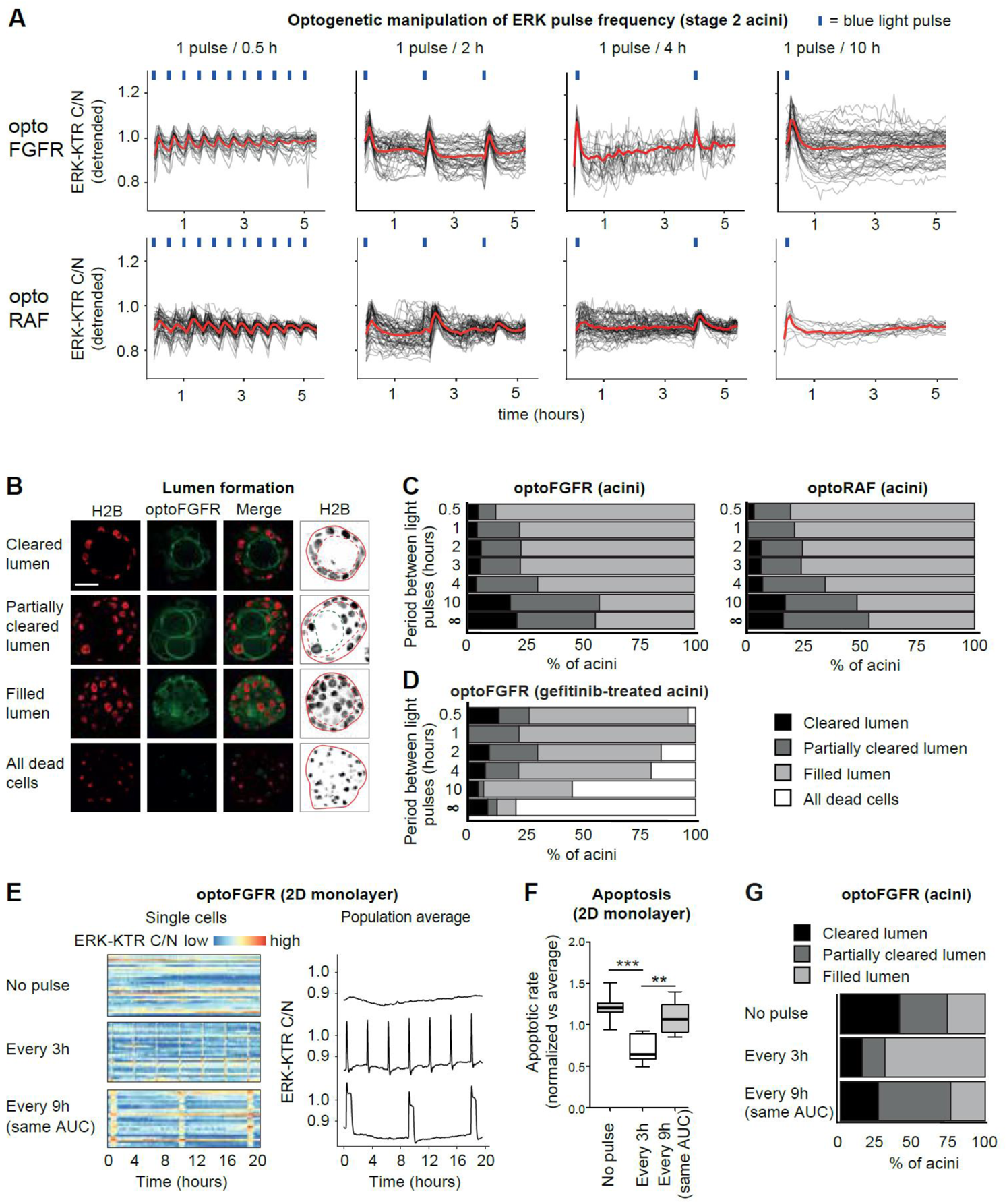
Controlling survival/apoptosis decisions with optogenetic actuators. (A) Overlayed detrended ERK activity trajectories from optoFGFR or optoRAF expressing acinar cells stimulated with different blue light pulse frequencies. (B) Colored: Equatorial optical sections of acini displaying cleared, partially cleared or filled luminal space or underwent complete cell death at stage 4. Black and white: maximal intensity projections of equatorial Z planes spanning 12 μm. Acini borders (red lines), luminal space (dashed red lines), and the border between the cleared and filled part of the luminal space (dashed green lines) are indicated. Scale bar = 20 μm. (C) Percentages of acini displaying cleared, partially cleared or filled luminal space at day 14, after 7 days on an LED plate that emitted blue light pulses at defined intervals. N = 36 - 72 acini per condition from 2 independent replicates. Pearson’s chi-squared test. optoFGFR: X^2^ (12 degrees of freedom, N = 384) = 78, P < 0.001. optoRAF: X^2^ (12 degrees of freedom, N = 326) = 32, P < 0.005 (D) Percentages of acini displaying cleared, partially cleared or filled luminal space or complete cell death at day 14, after 7 days in the presence of gefitinib on an LED plate that emitted blue light pulses at defined intervals. N = 24 - 46 acini per condition. X^2^ (15 degrees of freedom, N = 201) = 79, P < 0.001. (E) MCF10A cells in monolayer culture were stimulated every 3 hours with blue light pulses or every 9 hours with the same AUC, achieved with 20 consecutive blue light pulses, and compared with unstimulated cells. Randomly selected trajectories (left) and whole population average (right). (F) Distribution of apoptotic rates in 5 biological replicates each one normalized on the experiment mean. t-test (**, P < 0.01; ***, P < 0.001). (G) Percentages of optoFGFR-expressing acini that displayed cleared, partially cleared or filled luminal space at day 14, after 7 days on an LED plate that emitted either a single blue light pulse every 3 hours, 20 subsequent blue light pulses every 9 hours or no blue light pulses. N = 27-39 acini per condition. X^2^ (4 degrees of freedom, N = 104) = 23, P < .001.

Because endogenous ERK pulses occur on top of optogenetically-induced ones, we used optoFGFR to evoke different ERK pulse frequencies in EGFR-inhibited acini that do not exhibit spontaneous ERK pulses. Here, unstimulated acini and those stimulated every 10 hours displayed cell death. In marked contrast, ERK pulses applied at 0.5, 1, 2, 3 or 4 hours led to cell survival (Figure 5D). These results show that cells in acini must experience at least one ERK pulse every 4 hours to survive.

### ERK frequency but not integrated ERK activity regulates the survival fate

Next, we explored if the frequency of ERK pulses or the total integrated ERK activity over time controls survival. To test this, we used optoFGFR to induce synthetic ERK pulses of different widths, that when applied at different frequencies can evoke the same integrated ERK activity over a specific time period. Since ERK pulses display an identical shape and promote survival for 4 hours both in serum-deprived 2D monolayers (Gagliardi et al., 2021) and 3D acini, we first identified light stimulation schemes capable of inducing ERK pulses of different widths in monolayers. We applied a series of 1 to 86 successive blue light pulses delivered at 2-minute intervals, and recorded the resulting ERK dynamics (Figure S4C). We have previously shown that light stimulation applied at this frequency leads to sustained optoFGFR activity (Dessauges et al., 2021). The integrated ERK activity (area under the curve = AUC) displayed a linear relationship with the number of light pulses (Figure S4D). 20 blue light pulses delivered every 2 minutes led to a single ERK pulse with a three-fold increase in integrated ERK activity. We then applied 2 distinct optoFGFR stimulation schemes consisting of 1 AUC equivalent of ERK activity evoked every 3 hours versus 3 AUC equivalents of ERK activity evoked every 9 hours, resulting in the identical integrated ERK activity over a period of 9 hours (Figure 5E). We found that the first but not second optogenetic stimulation scheme induced cell survival in serum-deprived 2D cultures (Figure 5F). We then performed the identical experiment in stage 2 acini by applying the two optogenetic stimulation schemes using LITOS for 7 days. Quantification of lumen formation efficiency at day 14 revealed increased inner cell survival when the first, but not the second, optogenetic stimulation scheme was applied (Figure 5G). These results show that the frequency of ERK pulses rather than the integrated ERK activity regulates survival.

Finally, we also used our optogenetic toolkit to explore a number of scenarios of how different ERK frequencies in inner/outer cells might be regulated in stage 2 acini. These results are discussed in the Supplemental Text and Figure S5.

### Oncogenic PI3K signaling increases ERK frequency leading to loss of acinar lumen formation

We then explored ERK dynamics in the context of a pathological alteration of acinar morphogenesis induced by the H1047R mutation in the alpha subunit of PI3K (PIK3CA), which is frequently mutated in breast cancer (Cancer Genome Atlas Network, 2012). The H1047R PIK3CA mutation leads to absence of lumen in MCF10A knockin acini (Berglund et al., 2013; Chakrabarty et al., 2010; Chen et al., 2013, Isakoff et al., 2005; Lauring et al., 2010), as well as ductal hyperplasia in a transgenic mouse model (Tikoo et al., 2012). H1047R PIK3CA MCF10A knock-in cells have been shown to display elevated ERK activity using western blot (Gustin et al., 2009), strongly suggesting the existence of a crosstalk between oncogenic PI3K and MAPK/ERK signaling. Consistently, we found that H1047R PIK3CA MCF10A knock-in cells cultured as monolayers displayed higher median ERK frequency than their wild-type (WT) counterparts (Figure 6A-C), while maintaining similar ERK pulse shape, duration and amplitude (Figure 6D, S6A,B). The finding of a similar ERK pulse shape in WT and mutant cells indicates that the PI3K to MAPK crosstalk must occur upstream of Ras because the MAPK network structure that shapes ERK dynamics is maintained (Kholodenko et al., 2010). The increased ERK frequency suggests the involvement of receptor level interactions (Sparta et al., 2015). To better understand the effect of the PIK3CA H1047R on acinar morphogenesis, we evaluated proliferation and apoptosis during different stages (Figure 6E-I). Stage 1 mutant acini displayed increased proliferation compared to their WT counterparts, as evidenced by augmented cell numbers, and geminin quantification (Figure 6E-G). While remaining slightly higher than in WT acini, proliferation also diminished during stages 2 - 4, but displayed a small upshoot during stage 4 (Figure 6E,G). In contrast to the steep apoptosis rise observed starting on day 7 in WT acini, apoptotic rates remained lower in PIK3CA mutant acini at all stages (Figure 6E,H). Increased proliferation at stage 1, and decreased apoptosis at stages 3 and 4, thus mostly contribute to increased cell number and absence of lumen formation in PIK3CA-mutant acini (Figure 6I).

**Figure 6.**
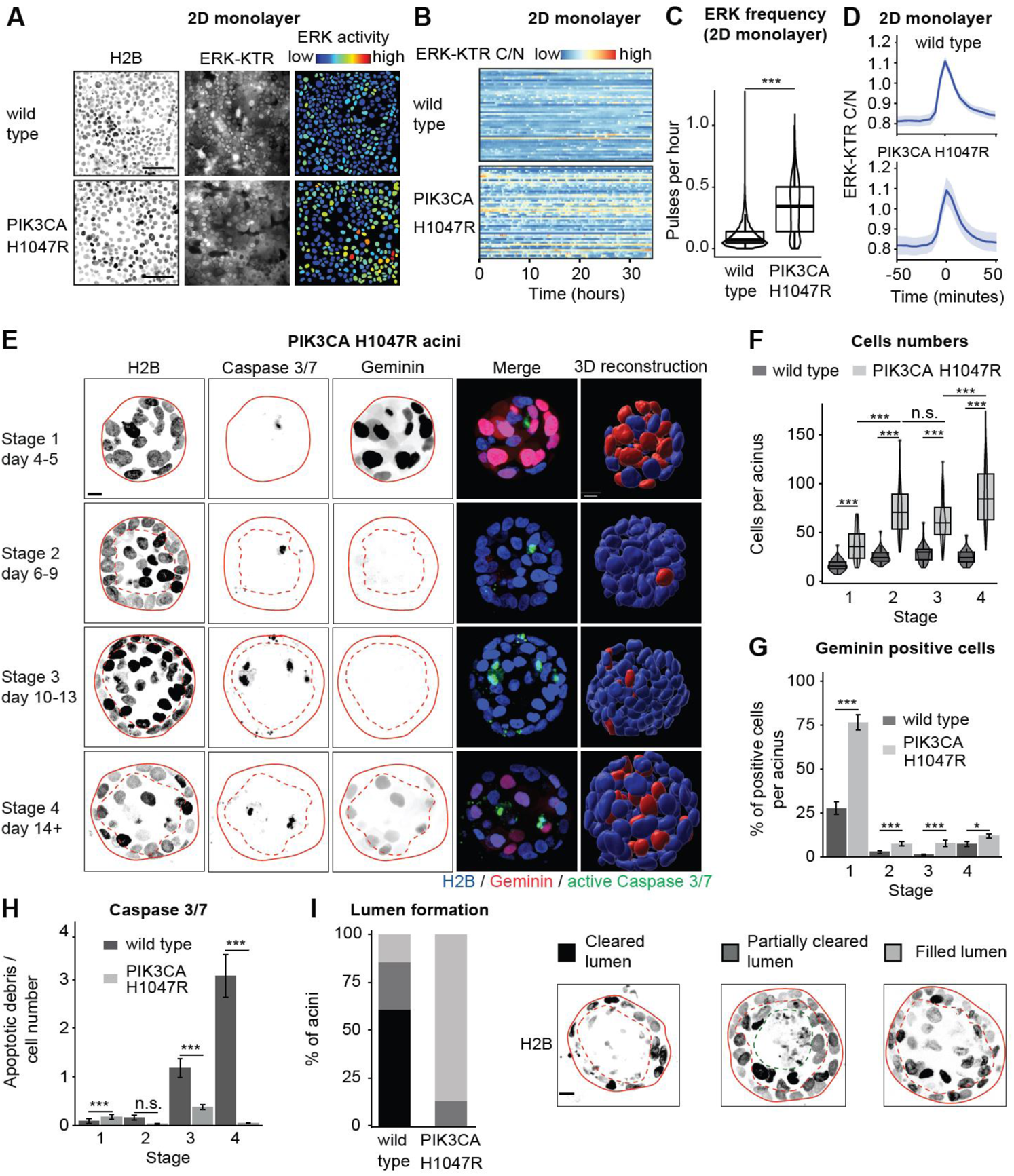
Increased 2D monolayer ERK frequency and altered acinar morphogenesis of PIK3CA H1047R cells. (A) Micrographs of WT and PIK3CA H1047R MCF10A 2D monolayers expressing fluorescent H2B (left) and ERK-KTR (middle). Right: nuclei of the same cells color-coded by ERK-KTR ratio. Scale bar = 100 μm. (B) Heatmap of single-cell ERK trajectories in WT and PIK3CA H1047R monolayers. (C) ERK frequencies in WT and PIK3CA H1047R monolayer cells. (D) Average ERK trajectories from isolated pulses in WT and PIK3CA H1047R cells within monolayers. 95% confidence intervals are shown. Time = 0 corresponds to maximal amplitude of peaks. (E) Micrographs and 3D reconstructions of H2B, caspase 3/7 fluorogenic substrate and geminin signals in PIK3CA H1047R acini at different stages. Micrographs show maximal intensity projections of equatorial Z planes spanning 12 μm. Plain lines mark the borders of the acini, dashed lines mark the outer cell layer. Scale bar = 10 μm. (F) Cell numbers per acinus at the different stages. N = 54 - 60 PIK3CA H1047R acini each. (G) Fraction of Geminin positive cells per acinus at different stages. (H) Number of Caspase 3/7 apoptotic debris divided by the acinar cell number at different days. (I) Percentages of acini that either displayed a cleared, partially cleared or filled luminal space at day 14. Pearson’s chi-squared test: X^2^ (2 degrees of freedom, N = 82) = 50, P < 0.001. Representative examples used for classification are shown (maximal intensity projections of equatorial Z planes spanning 12 μm). Acini borders (red lines), luminal space (dashed red lines), and the border between the cleared and filled part of the luminal space (dashed green lines) are indicated. Scale bar = 10 μm. (F-I) Measurements taken on the same acini. WT acini are the same as in figure S1, for comparison. (G, H) Error bars represent standard error of the mean. (C, F, G and H) Wilcoxon tests (n.s., P > 0.05; *, P < 0.05; **, P < 0.01; ***, P < 0.001).

We then evaluated ERK dynamics during the pathological acinar morphogenesis induced by PIK3CA H1047R. As in monolayers, PIK3CA-mutant acini displayed ERK pulses like those of WT acini (Figure 7A). Stage 1 rotating PIK3CA-mutant acini displayed ERK frequencies as high as those observed in their WT counterparts (Figure 7B). However, stage 1 non-rotating PIK3CA-mutant acini did not display the decreased ERK frequency observed in WT (Figure 7B). Stage 2 PIK3CA-mutant acini displayed increased ERK frequency compared to WT, as well as prominent ERK waves (Figure 7B-D, Movie S6). However, most likely due to their heterogeneity, we could not pinpoint a specific feature of ERK waves associated with the increased ERK frequency observed in PIK3CA-mutant versus WT acini. PIK3CA-mutant stage 2 displayed higher ERK frequencies than WT acini, leading the inner cells of PIK3CA-mutant to exhibit a similar ERK frequency than the outer cells of WT acini (Figure 7B,C). Amplitude and duration of ERK pulses were similar for both WT and PIK3CA H1047R acinar cells at all stages (Figure S6C,D).

**Figure 7.**
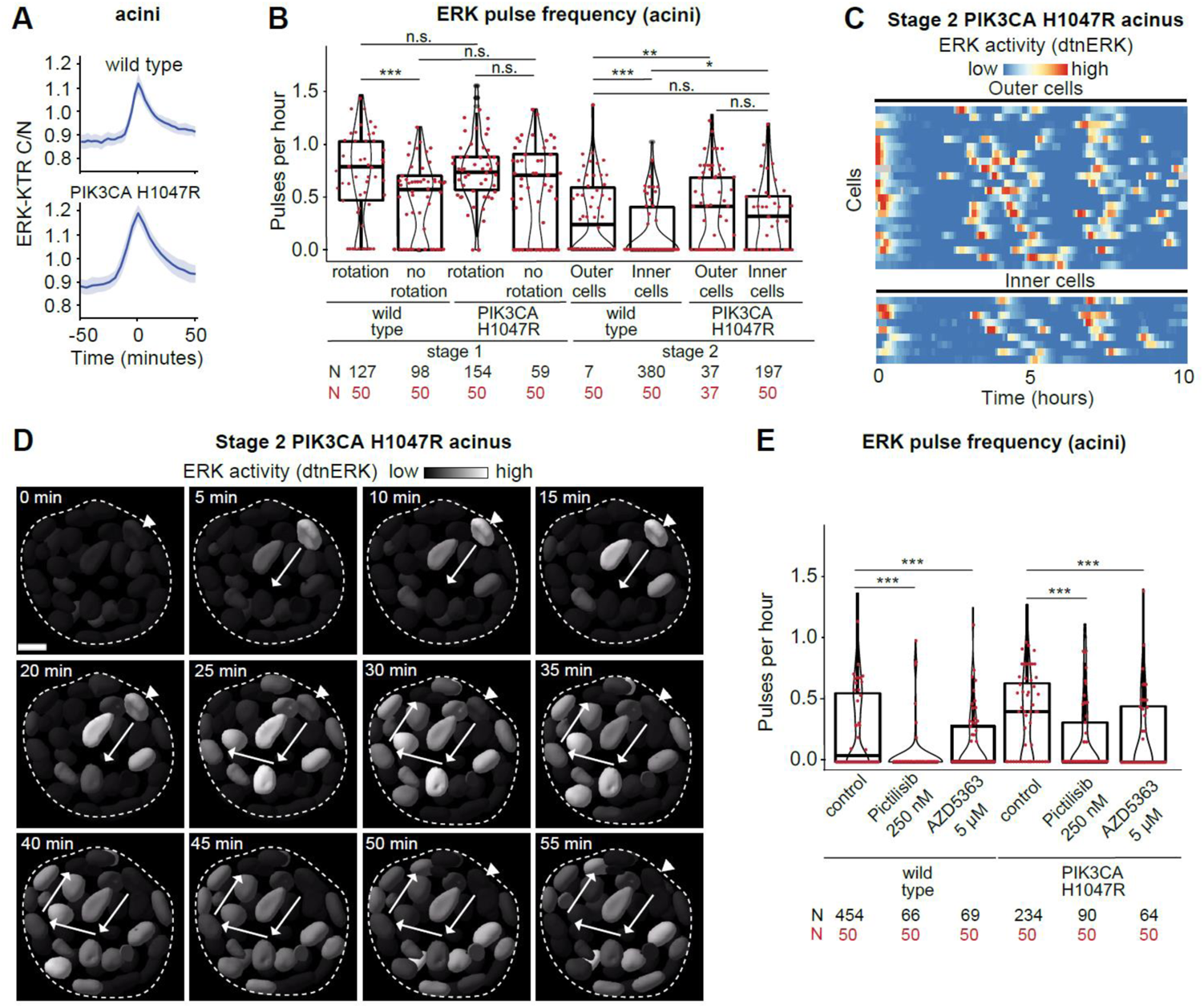
Increased ERK frequency in PIK3CA H1047R mutant acini. (A) Average ERK trajectories from isolated pulses in WT and PIK3CA H1047R cells within acini. 95% confidence intervals are shown. Time = 0 corresponds to maximal amplitude of peaks. (B) ERK frequencies of WT and PIK3CA H1047R cells at different stages and locations within the acinus. Mutant trajectories pooled from 7 (stage 1 rotation), 2 (stage 1 no rotation) and 6 (stage 2) acini. (C) Heatmap of detrended/normalized single-cell ERK trajectories in outer and inner cells of a representative stage 2 PIK3CA H1047R acinus. (D) Representative time-series of an ERK wave in a stage 2 PIK3CA H1047R acinus (dashed line) cross section. Nuclei are color-coded by ERK activity levels. Arrows show directionality of activation. Arrowhead indicates the initiator cell. Scale bar = 10 μm. (E) ERK frequencies of WT and PIK3CA H1047R cells from control acini and acini treated with 250 nM pictilisib or 5 µM AZD5363. (B and E) wt control data is the same as in figure 2 and 3, and shown again for comparison. Wilcoxon tests (n.s., P > 0.05; *, P < 0.05, **, P < 0.01; ***, P < 0.001).

Pictilisib-mediated PI3K or AZD5363-mediated Akt inhibition decreased ERK frequency in both WT and mutant acini (Figure 7E). Stimulation of WT acini with insulin-like growth factor (IGF1), that primarily activates PI3K-Akt signaling (Myers et al., 1993), also resulted in increased ERK pulse frequency (Figure S7A,B). Gefitinib-mediated EGFR inhibition abolished ERK pulses in PIK3CA-mutant acini, suggesting an EGFR-dependent mechanism (Figure S7C). Batimastat-mediated MMP inhibition in PIK3CA H1047R cells led to reduction of ERK phosphorylation to levels observed in WT cells, without affecting Akt signaling (Figure S7D). PIK3CA H1047R MCF10A cells have been shown to exhibit increased expression of the EGFR-ligand amphiregulin in comparison to WT cells (Young et al., 2015), possibly explaining the increase in EGFR-dependent ERK frequency. Consistently, we observed increased amphiregulin expression at mRNA level in PI3K-mutant when compared with WT cells. Amphiregulin expression levels were were decreased upon pictilisib-mediated inhibition of PI3K activity in PIK3CA mutant cells (Figure S7E). Together, these results strongly suggest that the PI3K to MAPK/ERK crosstalk that regulates ERK frequency involves amphiregulin/MMP-activation of EGFR. Increased survival in PIK3CA H1047R acini might therefore involve increased ERK frequency through this crosstalk mechanism, in addition to PI3K-Akt signaling.

## Discussion

Recent work in epithelial monolayers have revealed the existence of non-periodic single-cell ERK pulses, whose frequency controls apoptosis, survival or proliferation fates (Albeck et al., 2013; Aoki et al., 2013, Gagliardi et al., 2021; Valon et al., 2021). At the level of a cell population, these ERK pulses can be stochastic when cells are stimulated with EGF (Albeck et al., 2013), or can be organized as collective ERK waves during collective cell migration (Aoki et al., 2017; Hino et al., 2020), cancer cell extrusion (Aikin et al., 2020), or spatial regulation of survival in response to stress (Gagliardi et al., 2021; Valon et al., 2021). Here we show that similar ERK pulses/waves coordinate fate decisions during mammary acinar morphogenesis. As in the cell systems mentioned above, ERK pulses are triggered by MMP-mediated cleavage of pro-EGF ligands and subsequent activation of EGFR. Downstream of EGFR, MAPK network properties such as ultrasensitivity and negative feedback (Huang and Ferrell, 1996, Kholodenko et al., 2010) might allow to translate minute amounts of MMP-cleaved pro-EGF into clear cut digital ERK pulses. Consistently, the slightly lower amplitude of spontaneous ERK pulses versus those induced by acute EGF stimulation suggests that the EGFR/MAPK system functions at the threshold input to generate digital ERK pulses (Figure 1G). This might allow the MMP/EGFR/MAPK signaling network to translate small variations in the EGFR input into frequency-modulated regimes of ERK pulses that can specify proliferation, survival and apoptosis. Note that exogenous addition of EGF impedes apoptosis-mediated lumen formation (Gaiko-Shcherbak et al., 2015), further suggesting that small amounts of EGFR ligands synthesized by the acinus itself are necessary for self-organisation of its morphogenesis. This MMP/EGFR/MAPK network has also been shown to produce ERK trigger waves in which activated cells sequentially switch on ERK pulses in adjacent cells (Aoki et al., 2017; Boocock et al., 2020; Hino et al., 2020). Our results strongly suggest that the ERK waves observed in stage 2 acini rely on this mechanism. Thus, a relatively simple signaling network might allow to produce conserved and sophisticated ERK behaviors in epithelial collectives. This excitable MMP/EGFR/MAPK network that generates pulsatile ERK activity strongly contrasts with the oscillatory ERK behavior observed in the segmentation clock in vertebrate embryos that is regulated on slower timescales by rhythmic transcriptional regulation of MAPK phosphatases (Hubaud and Pourquié, 2014). We speculate that the pulsatile MAPK network observed in epithelia provides an opportunity to constantly sense and react to environmental inputs such as growth factors and insults to warrant epithelial homeostasis during acinar development. This illustrates how the MAPK network can be differently wired to produce distinct ERK dynamics at different timescales depending on the developmental context.

### ERK dynamics regulate collective migration and proliferation in stage 1 acini

We found that stage 1 acini displayed high frequency, asynchronous ERK pulses during rapid collective cell migration that leads to a global rotation behavior of the acinus (Figure 2). This rotation behavior has previously been implicated in the morphogenesis of spherical tissue buds during mammary organogenesis (Fernández et al., 2021). During migration of 2D epithelial sheets, ERK waves co-ordinate myosin activity necessary for collective motility, and are shaped by a mechanochemical feedback from myosin to ERK (Aoki et al., 2017; Boocock et al., 2020; Hino et al., 2020). Further, ERK also has been shown to control myosin activity in MCF10A acini single-cell motility (Pearson and Hunter, 2007). We therefore propose that asynchronous ERK pulses spatially coordinate myosin contractility necessary for this collective motility behavior. This is consistent with our result that optogenetic synchronization of ERK pulses immediately leads to decreased collective motility (Figure 4C-F). The transition to a state of slower motility in late stage 1 acini has been shown to result from assembly of an endogenous laminin-rich basement membrane (Wang et al., 2013). We speculate that assembly of this basement membrane might modify the myosin-ERK mechanochemical feedback loop mentioned above, leading to decreased ERK frequency and motility, allowing to regulate the transition from proliferation to quiescence. Assembly of the basement membrane might therefore act as a checkpoint coordinating ERK frequency-dependent regulation of motility and transition from proliferation to quiescence. Future experimental/modeling studies will be necessary to further refine this hypothesis.

### ERK waves spatially regulate apoptosis and survival fates during stage 2 acinar lumen morphogenesis

Our experiments in stage 2 acini suggest that spatial control of different ERK frequencies regulates survival in outer versus apoptosis fates in inner acinar cells. Outer cells display a median ERK frequency of one pulse every 4 hours, while inner cells display lower ERK frequencies (Figure 3A). This is consistent with the ability of one ERK pulse to provide about 4 hours of survival in MCF10A monolayers (Gagliardi et al., 2021). Using optogenetic control of ERK pulses, we excluded mechanisms such as differential growth factor receptor sensitivity or refractory time of inner and outer cells for regulation of different ERK frequencies (Supplementary Text, Figure S5A,B). Instead, our results suggest a role for collective ERK waves to define the outer and inner spatial domains of high and low ERK frequency, that respectively specify survival and apoptosis fates (Figure 3B-I). ERK wave properties such as that they are initiated mostly in the outer layer, and propagate more efficiently in the outer versus the inner layer, might dynamically specify the two domains of ERK frequencies on timescale of hours throughout the 7 days of the acinus cavitation process. While apoptosis is predominantly responsible for the clearance of luminal cells in acini (Debnath et al., 2002), we cannot exclude that an alternative mechanism such as autophagy resulting from metabolic defects in inner cells, which is regulated by EGFR - PI3K signaling might also contribute to this process (Schafer et al., 2009).

### ERK pulse frequency but not integrated activity regulates survival versus apoptosis fates

To formally test if ERK frequency regulates survival/apoptosis fates, we used optogenetic stimulation of all the cells of acini to causally link ERK frequency with lumen formation. When used to evoke ERK pulses for up to at least every 4 hours, both optoFGFR and optoRaf led to loss of lumen formation (Figure 5C). ERK pulses evoked every 10 hours were not sufficient to rescue apoptosis. These experiments also functioned when EGFR was completely inhibited (Figure 5D). By optogenetically varying the ERK pulse width, we also showed that ERK frequency rather than the integrated ERK activity over time regulates survival versus apoptosis fates, both in monolayers and in stage 2 acinus lumen formation (Figure 5E-G). Our data therefore suggests that short ERK pulses are the signaling unit that allows cells in acini to commit to survival for about 4 hours. This might allow cells to integrate signaling inputs that fluctuate on timescales of minutes/hours, to dynamically control a morphogenetic program on timescales of days. Interpretation of a specific ERK frequency into survival/apoptosis fates might involve the ERK substrate Bcl-2-like protein 11 (Harada et al., 2004) or ERK-dependent transcriptional control of immediate early genes (IEGs) (Avraham and Yarden, 2011). IEGs produce transcripts with lifetimes of around 30 minutes, that encode proteins with lifetimes of 1-3 hours. Notably, IEGs include Jun and Fos transcription factors which are important regulators of cell survival (Shaulian and Karin, 2001). IEG’s short lifetime is compatible with the ERK frequency required for survival. Higher ERK frequencies as observed in stage 1 acini might control proliferation by regulation of IEGs such as Fra-1 (Gillies et al., 2017).

### Oncogenic PI3K signaling modulates ERK frequency contributing to aberrant acinar morphogenesis

Our results show that the breast cancer relevant PIK3CA H1047R mutation increases ERK frequency, which might contribute at least in part to increased proliferation and survival leading to larger acini without lumen formation, that phenocopies ductal hyperplasia observed in mouse models (Tikoo et al., 2012). We show that this crosstalk from PI3K to ERK signaling feeds into the control of ERK frequency both in WT and PIK3CA H1047R monolayers and acini (Figure 7, S6-7). This crosstalk also functions downstream of IGF1, that primarily activates PI3K-Akt signaling (Myers et al., 1993) (Figure S7A,B), suggesting that it is a conserved feature downstream of multiple receptor tyrosine kinases. During stage 1, PIK3CA H1047R acini display similarly high ERK frequencies as in WT acini (Figure 7B). During stage 2, PIK3CA H1047R acini then still display a substantial decrease in ERK frequency as in WT acini compared to stage 1 (Figure 7B), which correlates with decreased proliferation (Figure 6E-H). However, ERK frequencies remain slightly higher both in outer and inner cells in PIK3CA H1047R versus WT acini, correlating with the strong survival phenotype in inner cells, and absence of lumen formation. These results suggest that the control of ERK frequency is still subject to some degree of regulation even when the crosstalk from PIK3CA H1047R is constitutively switched on. Because the ERK pulse shape is identical in WT and PIK3CA mutant cells, this crosstalk must occur upstream of the core Raf/MEK/ERK circuit. Consistently, ERK activation is sensitive to both EGFR and MMP inhibition in PIK3CA H1047R cells (Figure S7C,D). As reported previously (Young et al., 2015), our data strongly suggests that constitutive PI3K activity in PIK3CA H1047R cells leads to increased expression of the EGFR ligand amphiregulin that in turn might increase ERK frequency (Sternlicht et al., 2005) (Figure S7E). PIK3CA H1047R also has been found to decrease expression of the protein tyrosine phosphatase receptor type F (PTPRF) (Young et al., 2015), which might further augment EGFR excitability and thus ERK frequency. Further work is required to elucidate the specific contributions of PI3K and ERK signaling to control the survival/apoptosis fate decisions and how the PI3K/ERK crosstalk might spatially fine tune ERK frequency. Our result strongly suggests that oncogenic PI3K signaling-induced aberrant spatial regulation of ERK frequency contributes to pathological acinar morphogenesis.

## Conclusion

We provide an initial characterization into how single-cell ERK dynamics control fate decisions in space and time during the morphogenesis of a simple prototype organ structure. Future studies are required to mechanistically understand how the different dynamic signaling states are encoded and spatially organized, and if they provide robustness against environmental perturbations occurring during development. This will require the ever-expanding arsenal of optogenetic tools to manipulate specific cells and evaluate how the cell collective responds. Further questions include how additional signaling pathways might fine tune this morphogenetic process, and how the ERK frequency is decoded into transcriptional programs that actuate the different fates that shape acinus morphogenesis.

## Material and methods

### 2D cell culture

MCF10A cells were cultured in DMEM/F12 supplemented with 5% horse serum, 20 ng/ml recombinant human EGF (Peprotech), 0.5 μg/ml hydrocortisone (Sigma-Aldrich/Merck), 10 μg/ml insulin (Sigma-Aldrich/Merck), 200 U/ml penicillin and 200 μg/ml streptomycin. The PIK3CA H1047R knockin cell line (Gustin et al., 2009) was a gift of Ben Ho Park. We regularly verified the presence of the mutation by sequencing the corresponding genomic locus. To generate stable cell lines, cells were transfected with FuGene (Promega) according to the manufacturer’s protocol and clones were selected by antibiotic resistance and image-based screening.

### 3D cell culture

For acinus formation, single MCF10A cell suspensions were mixed with 4 volumes of growth factor-reduced Matrigel (Corning) at 4 °C and spread evenly on the surface of glass bottom cell culture plates at a concentration of 1.5 x 10^4^ cells/cm^2^. Acini were cultured in DMEM/F12 supplemented with 2% horse serum, 20 ng/ml recombinant human EGF, 0.5 μg/ml hydrocortisone, 10 μg/ml insulin, 200 U/ml penicillin and 200 μg/ml streptomycin. Horse serum, insulin and EGF were removed after 3 days of culture. For live imaging, 25 mM Hepes was added to the medium prior to mounting on the microscope. CellEvent Caspase 3/7 Green Detection Reagent was obtained from Thermo Fisher Scientific and used according to the manufacturer’s protocol.

### Plasmids

ERK-KTR-mTurquoise2 and ERK-KTR-mRuby2 were generated by fusion of the coding region of ERK-KTR (Regot et al., 2014) with that of mTurquoise2 (Goedhart et al., 2012) or mRuby2 (Lam et al., 2012). H2B-miRFP703 was generated by fusion of the coding region of human H2B clustered histone 11 (H2BC11) with that of miRFP703 (Shcherbakova et al., 2016). Geminin-mCherry was generated by fusion of the ubiquitylation domain of human Geminin (Sakaue-Sawano et al., 2017) to mCherry. The above mentioned fusion proteins were cloned in the piggyBac vectors pMP-PB, pSB-HPB (Balasubramanian et al., 2016) (gift of David Hacker, EPFL), or pPB3.0.Blast, an improved version of pPB generated in our lab.

pPB3.0-PuroCRY2-cRAF-mCitrine-P2A-CIBN-KrasCT, referred to in the manuscript as OptoRAF, was generated in the following way: The CRY2-cRaf sequence was excised from pCX4puro-CRY2-cRAF (gift from Kazuhiro Aoki, (Aoki et al., 2017)) using EcoRI and NotI. mCitrine was PCR amplified from the optoFGFR plasmid, while adding NotI and XhoI sites, and digested. Both sequences were ligated into pPB3.0-Puro, previously digested with EcoRI and XhoI. The GSGP2A-CIBN-KRasCT sequence (synthesized by GENEWIZ) was digested with BsrGI and AflII and ligated into pPB3.0-Puro-CRY2-cRAF-mCitrine.

The piggyBac plasmids were co-transfected with a helper plasmid expressing a hyperactive piggyBac transposase (Yusa et al., 2011).

Lyn-cytoFGFR1-PHR-mCit, expressing myristoylated FGFR1 cytoplasmic region fused with the PHR domain of cryptochrome2 and mCitrine (gift from Won Do Heo (Addgene plasmid # 59776), (Kim et al., 2014)), referred to in the manuscript as OptoFGFR, was subcloned in a lentiviral backbone for stable cell line generation.

### Imaging

All acini images were acquired on an epifluorescence Eclipse Ti2 inverted fluorescence microscope (Nikon) equipped with a CSU-W1 spinning disk confocal system (Yokogawa) and a Plan Apo VC 60X water immersion objective (NA = 1.2). For time lapse imaging, laser-based autofocus was used. Images were acquired with a Prime 95B or a Prime BSI sCMOS camera (both Teledyne Photometrics) at 16-bit depth. Temperature, CO2 and humidity were controlled throughout live imaging with a temperature control system and gas mixer (both Life Imaging Services).

All monolayer cell images were acquired on an epifluorescence Eclipse Ti inverted fluorescence microscope (Nikon) with a Plan Apo 20x air objective (NA = 0.8) or a Plan Apo 40X air objective (NA = 0.9). Laser-based autofocus was used throughout imaging. Images were acquired with an Andor Zyla 4.2 plus camera at 16-bit depth. Both microscopes were controlled by NIS elements (Nikon).

### Inhibitors and growth factors

Gefitinib was obtained from Sigma-Aldrich/Merck and used at a concentration of 10 μM. Trametinib was obtained from Selleck Chemicals and used at a concentration of 5 μM. Batimastat was obtained from MedChem Express and used at a concentration of 30 μM. Pictilisib was obtained from Selleck Chemicals and used at a concentration of 250 nM. AZD5363 was obtained from Selleck Chemicals and used at a concentration of 5 μM. IGF1 was obtained from Peprotech and used at a concentration of 100 nM.

### Optogenetic experiments

For short term optogenetics experiments performed directly on the microscope, acini were illuminated with wide field blue light (470 nm LED) at defined time points during spinning disc time lapse imaging. Acini expressing optoFGF were illuminated for 100 ms at 50% LED intensity. Acini expressing optoRAF were illuminated for 100 ms at 60% LED intensity. The NIS elements JOBS module was used to program the imaging and stimulation patterns.

The 2D monolayer optogenetic experiments were executed by culturing MCF10A expressing optoFGFR, ERK-KTR-mRuby2 and H2B-miRFP703 as confluent monolayer on 24 well plates with glass bottom. Optogenetic stimulation was done with 488 nm LED light at 100% light intensity for 100ms and using a 20x air objective. To generate larger ERK activity pulses, cells were stimulated with trains of 488 nm light pulses with a 2 min interval. The area under the curve (AUC) of ERK activity was calculated using the ERK activity levels before and after the ERK pulses to set the baseline. The correlation function between the number of 488 nm light pulses and AUC was obtained by linear regression. The effects of different frequencies with same AUC on apoptosis was measured after 24 hours of time-lapse acquisition and manual annotation of apoptotic events on the base of morphological alterations of cell nucleus. For long term optogenetic stimulation with the LITOS system, glass bottom 96-well cell culture plates with 7-day old acini were fitted on a 32 x 64 RGB LED matrix (Boxtec) inside a cell culture incubator. The matrix was connected to a custom printed circuit board with an ESP32 microcontroller. This system was programmed to emit 1-minute blue light pulses at maximal intensity at defined intervals for 7 days, after which lumen formation efficiency was assessed. Acini were fixed with 4% paraformaldehyde prior to imaging.

### Image analysis of 3D acini

The open source LEVERJS software (Cohen, 2014; Wait et al., 2014; Winter et al., 2016) was used to analyze the 3D time lapse movies. The LEVERJS software was updated to include improved processing and visualization capabilities. The processing pipeline began with a GPU-accelerated 3D non-local means denoising algorithm (Wait et al., 2019). After denoising, a new ensemble-based segmentation algorithm was applied. This ensemble segmentation combined an adaptive thresholding into foreground/background regions with an anisotropic 3D Laplacian of Gaussian filter targeted to a specific cell radius to separate touching cells (Winter et al., 2019). The base segmentation was run at different cell radii and the results were combined using unsupervised learning techniques from the field of algorithmic information theory (Cohen et al., 2009). Here, the radii evaluated ranged from 2.5 μm to 4 μm in 0.5 μm steps. These values were set empirically based on expected cell size ranges. Following segmentation, the cells were tracked using Multitemporal Association Tracking (Winter et al., 2011); (Winter et al., 2012).

Following segmentation and tracking of the image sequences, the ERK-KTR signal was extracted and processed to a detrended and normalized signal. To extract the ERK-KTR signal, distance transforms were computed for each segmented image. The interior distance transform assigned each cell interior voxel a numeric value indicating its distance starting at the cell boundary and increasing to the centroid. The exterior distance transform assigned each boundary voxel a numeric value indicating its distance to the nearest cell-assigned voxel. The exterior distance transform also provided the identity of the nearest cell-assigned voxel for each background voxel. The ERK-KTR signal was computed as the ratio of the image values around the center of the cell to the image values around the boundary of the cell. The center region of the cell included voxels in the 95^th^ percentile of the interior distance transform. For the boundary region of the cell we included interior voxels within one unit of the boundary and exterior voxels within three units of the boundary. The resulting ERK-KTR signal was computed as the ratio of the median voxel value in the outer region to the median voxel value in the inner region. The extracted ERK-KTR signals for each cell had different base intensities and showed different amounts of fluctuation. To normalize this and to allow for quantitative comparison and visual representation with different cell ERK-KTR expression levels we computed a detrended and normalized ERK-KTR signal as follows. The signal S was first detrended by subtracting the median filter signal, *S_d_* = *S* - median_filter(*S*). The signal was then normalized to the range [0,1] using *S_nd_* = *S_d_* / max(*S_d_*), unless the signal range in the detrended trajectory was below an empirically set threshold. Inner and outer cells in stage 2 acini were identified visually based on the 3D reconstructions in LEVERJS.

For segmentation and quantification of steady state Z-stacks, CellProfiler 3.1.8 (McQuin et al., 2018) with 3D functionalities was used. Nuclei were identified based on H2B-miRFP signals. Apoptotic debris were identified using Caspase 3/7 Green Detection kit signals using adaptive thresholding and watershed segmentation. Geminin-mCherry intensities were measured within nuclei voxel masks. ERK-KTR cytosolic/nuclear intensity ratios were generated by measuring median ERK-KTR intensities in the nuclear area and in a spherical voxel mask 1 pixel around the nuclear objects. Imaris software (Bitplane) was used for 3D rendering of confocal stacks and to track and measure motility parameters in Figures 2 and 4.

### Image analysis of 2D cell cultures

Nuclei of monolayer cells were segmented using Ilastik (Berg et al., 2019) and CellProfiler 3.0. Ilastik was used for training a random forest classifier on different pixel features of the H2B-miRFP channel and background pixels. The training data was generated by manual annotation of 20 - 50 cells. The resulting probability map was imported into CellProfiler for thresholding and segmentation. Cytosolic ERK-KTR fluorescence intensities were measured by expansion of the nuclear objects. Cells were tracked using μ-track 2.2.1 (Jaqaman et al., 2008).

### Data analysis

All analysis and visualization of ERK activity peaks and time series was performed with custom R/Matlab/Python code.

### ERK activity pulse detection

ERK activity peaks were identified and counted by the following steps. 1. Application of a short median filter to smooth the time series. 2. Application of a long median filter to produce a bias estimate which was subtracted from the smoothed time series. 3. Detrended time-series with real peaks were then identified by selecting those with an activity difference above an empirical threshold. Those were rescaled to [0,1] and a local maxima detection algorithm was used to identify peaks above an amplitude of 0.1.

### Identification of collective events

To identify waves of collective ERK activation we developed a custom code and implemented it in R. The algorithm works on a binarised signal that is calculated by detrending and normalising ERK-KTR cytosolic/nuclear intensity ratio time series as described above (Figure S3F,G). The algorithm searches for the first occurrence of cells with ERK switched on (Figure S3H). If several such cells exist and they are located within a threshold radius, they initiate the first collective event. A single active cell can also become a collective event. In the next time point, the algorithm repeats the search for active cells and compares their distances to cells in the previous frame. If new active cells are located within the threshold distance to active cells at the previous time point, they are added to respective collective events. If new active cells are located outside of the threshold distance, they form new collective events. This process of growing clusters of collective activity is repeated for all remaining time points. The resulting statistics include the total number of cells involved in a collective event, the duration and the average size of an event and the location (inner or outer layer) of the cells that initiates the event.

### Immunoblotting and qPCR

Cells were plated into 6-well dishes (2 x 10^5^ cells/well) and cultured for 48h. The resulting cell monolayers were washed twice with room temperature PBS, then starving medium was added. For immunoblotting 1uL/mL DMSO or 10ug/mL Batimastat was added together with the starving medium and cells were further cultured for 72h. Media was removed, monolayers were washed twice with ice-cold PBS, whole cell lysates were prepared and analyzed by immunoblotting as described before (Dessauges et al., 2021). Primary antibodies against the following proteins/epitopes were used: phospho-AKT^Ser473^ (cat. # 4058), AKT (cat. # 9272), phospho-p44/42 MAPK (Erk1/2)^Thr202/Tyr204^ (cat. # 4370), p44/42 MAPK (Erk1/2) (cat. # 4695), all from Cell Signaling Technologies, BioConcept Ltd. Secondary IRDye680LT- or IRDye800CW-conjugated anti-species specific IgGs were from LI-COR. For qPCR cells were first starved for 24h, then fresh starving medium containing 1uL/mL DMSO or 2uM and 10uM pictilisib was added and cells were cultured for 24h. Media was removed, monolayers were washed twice with ice-cold PBS, RNA was isolated using TRIzol reagent. Reverse transcription was done with the ProtoScript II reverse transcriptase kit (Bioconcept, M0368L). Real-time qPCR reactions were run using the MESA Green qPCR MasterMix Plus for SYBR Green assay (Eurogenetec, RT-SY2X-03+WOU) on the Rotor-Gen Q device (Qiagen). Each sample was tested in triplicate. Expression of the gene of interest was calculated using the 2^-ΔΔCt^ method.

The sequences of the primer sets used are as follows: AREG, 5’-ACA TTT CCA TTC TCT TGT CG-3’ (forward), and 5’-ACA TTT CCA TTC TCT TGT CG-3’ (reverse); FLJ22101, 5’-TTC CCT GTG GCA CTT GAC ATT-3’ (forward), and 5’-CTT TTG CCT CTG GCA GTA CTC A-3’ (reverse).

### Quantification and statistical analysis

All graphs were assembled and statistics were performed using R or Excel. Box plots depict the median and the 25th and 75th percentiles; whiskers correspond to minimum and maximum non-outlier values in Figures 1G, 2D, 3A, 3H, 3I, 5F, 6C, 6F, 7B, 7E, S1C, S3B, S3C, S3D, S3E, S4B, S6A, S6B, S6C, S6D and S7A. Dot plots show distribution of 50 randomly selected data points per condition, or all data points if there are less than 50. Red lines in ERK activity trajectories represent the population average. The statistical tests used and the significance thresholds are indicated in each respective legend.

## Data and code availability

The open-source code for LEVERJS is available at https://leverjs.net/git. All 3D datasets that were analyzed with LEVERJS for this publication can be browsed interactively at https://leverjs.net/mcf10a_3d.

## Acknowledgments

The authors are grateful to Ben Ho Park for providing the MCF10A PIK3CA H1047R knockin MCF10A cells, to Kazuhiro Aoki for the optoRaf construct, and to Won Do Heo for the optoFGFR construct. This work was supported by grants from the Swiss National Science Foundation and the Swiss Cancer Research Foundation to Olivier Pertz and from a Human Frontiers Science Program grant to Olivier Pertz and Andrew Cohen. We acknowledge support of the Microscopy Imaging Center of the University of Bern (https://www.mic.unibe.ch/).

## Author contribution

PE, PAG and OP designed the study. PE, PAG and AF performed experiments and analyzed data. MD and M-AJ analyzed data. CD provided expertise with the optogenetic tools. TH provided expertise with LITOS. ARC performed image analysis using LEVERJS. PE, PAG and OP wrote the paper.

## References

Aikin, T.J., Peterson, A.F., Pokrass, M.J., Clark, H.R., and Regot, S. (2020). MAPK activity dynamics regulate non-cell autonomous effects of oncogene expression. ELife 9.

Albeck, J.G., Mills, G.B., and Brugge, J.S. (2013). Frequency-modulated pulses of ERK activity transmit quantitative proliferation signals. Mol. Cell 49, 249–261.

Anderson, L.R., Sutherland, R.L., and Butt, A.J. (2010). BAG-1 overexpression attenuates luminal apoptosis in MCF-10A mammary epithelial cells through enhanced RAF-1 activation. Oncogene 29, 527–538.

Aoki, K., Kumagai, Y., Sakurai, A., Komatsu, N., Fujita, Y., Shionyu, C., and Matsuda, M. (2013). Stochastic ERK activation induced by noise and cell-to-cell propagation regulates cell density-dependent proliferation. Mol. Cell 52, 529–540.

Aoki, K., Kondo, Y., Naoki, H., Hiratsuka, T., Itoh, R.E., and Matsuda, M. (2017). Propagating wave of ERK activation orients collective cell migration. Dev. Cell 43, 305–317.e5.

Avraham, R., and Yarden, Y. (2011). Feedback regulation of EGFR signalling: decision making by early and delayed loops. Nat. Rev. Mol. Cell Biol. 12, 104–117.

Balasubramanian, S., Wurm, F.M., and Hacker, D.L. (2016). Multigene expression in stable CHO cell pools generated with the piggyBac transposon system. Biotechnol. Prog. 32, 1308–1317.

Berglund, F.M., Weerasinghe, N.R., Davidson, L., Lim, J.C., Eickholt, B.J., and Leslie, N.R. (2013). Disruption of epithelial architecture caused by loss of PTEN or by oncogenic mutant p110α/PIK3CA but not by HER2 or mutant AKT1. Oncogene 32, 4417–4426.

Berg, S., Kutra, D., Kroeger, T., Straehle, C.N., Kausler, B.X., Haubold, C., Schiegg, M., Ales, J., Beier, T., Rudy, M., et al. (2019). ilastik: interactive machine learning for (bio)image analysis. Nat. Methods 16, 1226–1232.

Boocock, D., Hino, N., Ruzickova, N., Hirashima, T., and Hannezo, E. (2020). Theory of mechanochemical patterning and optimal migration in cell monolayers. Nat. Phys.

Cancer Genome Atlas Network (2012). Comprehensive molecular portraits of human breast tumours. Nature 490, 61–70.

Chakrabarty, A., Rexer, B.N., Wang, S.E., Cook, R.S., Engelman, J.A., and Arteaga, C.L. (2010). H1047R phosphatidylinositol 3-kinase mutant enhances HER2-mediated transformation by heregulin production and activation of HER3. Oncogene 29, 5193–5203.

Chen, N., Eritja, N., Lock, R., and Debnath, J. (2013). Autophagy restricts proliferation driven by oncogenic phosphatidylinositol 3-kinase in three-dimensional culture. Oncogene 32, 2543–2554.

Ciarloni, L., Mallepell, S., and Brisken, C. (2007). Amphiregulin is an essential mediator of estrogen receptor alpha function in mammary gland development. Proc Natl Acad Sci USA 104, 5455–5460.

Cohen, A.R. (2014). Extracting meaning from biological imaging data. Mol. Biol. Cell 25, 3470– 3473.

Cohen, A.R., Bjornsson, C.S., Temple, S., Banker, G., and Roysam, B. (2009). Automatic summarization of changes in biological image sequences using algorithmic information theory. IEEE Trans. Pattern Anal. Mach. Intell. 31, 1386–1403.

Debnath, J., Mills, K.R., Collins, N.L., Reginato, M.J., Muthuswamy, S.K., and Brugge, J.S. (2002). The role of apoptosis in creating and maintaining luminal space within normal and oncogene-expressing mammary acini. Cell 111, 29–40.

Debnath, J., Muthuswamy, S.K., and Brugge, J.S. (2003). Morphogenesis and oncogenesis of MCF-10A mammary epithelial acini grown in three-dimensional basement membrane cultures. Methods 30, 256–268.

Dessauges, C., Mikelson, J., Dobrzynski, M., Jacques, M.-A., Frismantiene, A., Gagliardi, P.A., Khammash, M., and Pertz, O. (2021). Optogenetic actuator/biosensor circuits for large-scale interrogation of ERK dynamics identify sources of MAPK signaling robustness. BioRxiv.

Du, W.W., Fang, L., Li, M., Yang, X., Liang, Y., Peng, C., Qian, W., O’Malley, Y.Q., Askeland, R.W., Sugg, S.L., et al. (2013). MicroRNA miR-24 enhances tumor invasion and metastasis by targeting PTPN9 and PTPRF to promote EGF signaling. J. Cell Sci. 126, 1440–1453.

Fernández, P.A., Buchmann, B., Goychuk, A., Engelbrecht, L.K., Raich, M.K., Scheel, C.H., Frey, E., and Bausch, A.R. (2021). Surface-tension-induced budding drives alveologenesis in human mammary gland organoids. Nat. Phys. 17, 1130–1136.

Finlay, D., Healy, V., Furlong, F., O’Connell, F.C., Keon, N.K., and Martin, F. (2000). MAP kinase pathway signalling is essential for extracellular matrix determined mammary epithelial cell survival. Cell Death Differ. 7, 302–313.

Gagliardi, P.A., Dobrzyński, M., Jacques, M.-A., Dessauges, C., Ender, P., Blum, Y., Hughes, R.M., Cohen, A.R., and Pertz, O. (2021). Collective ERK/Akt activity waves orchestrate epithelial homeostasis by driving apoptosis-induced survival. Dev. Cell 56, 1712–1726.e6.

Gaiko-Shcherbak, A., Fabris, G., Dreissen, G., Merkel, R., Hoffmann, B., and Noetzel, E. (2015). The acinar cage: basement membranes determine molecule exchange and mechanical stability of human breast cell acini. PLoS ONE 10, e0145174.

Gillies, T.E., Pargett, M., Minguet, M., Davies, A.E., and Albeck, J.G. (2017). Linear Integration of ERK Activity Predominates over Persistence Detection in Fra-1 Regulation. Cell Syst. 5, 549–563.e5.

Goedhart, J., von Stetten, D., Noirclerc-Savoye, M., Lelimousin, M., Joosen, L., Hink, M.A., van Weeren, L., Gadella, T.W.J., and Royant, A. (2012). Structure-guided evolution of cyan fluorescent proteins towards a quantum yield of 93%. Nat. Commun. 3, 751.

Goglia, A.G., Wilson, M.Z., Jena, S.G., Silbert, J., Basta, L.P., Devenport, D., and Toettcher, J.E. (2020). A Live-Cell Screen for Altered Erk Dynamics Reveals Principles of Proliferative Control. Cell Syst. 10, 240–253.e6.

Gustin, J.P., Karakas, B., Weiss, M.B., Abukhdeir, A.M., Lauring, J., Garay, J.P., Cosgrove, D., Tamaki, A., Konishi, H., Konishi, Y., et al. (2009). Knockin of mutant PIK3CA activates multiple oncogenic pathways. Proc Natl Acad Sci USA 106, 2835–2840.

Harada, H., Quearry, B., Ruiz-Vela, A., and Korsmeyer, S.J. (2004). Survival factor-induced extracellular signal-regulated kinase phosphorylates BIM, inhibiting its association with BAX and proapoptotic activity. Proc Natl Acad Sci USA 101, 15313–15317.

Hino, N., Rossetti, L., Marín-Llauradó, A., Aoki, K., Trepat, X., Matsuda, M., and Hirashima, T. (2020). ERK-Mediated Mechanochemical Waves Direct Collective Cell Polarization. Dev. Cell 53, 646–660.e8.

Hiratsuka, T., Fujita, Y., Naoki, H., Aoki, K., Kamioka, Y., and Matsuda, M. (2015). Intercellular propagation of extracellular signal-regulated kinase activation revealed by in vivo imaging of mouse skin. ELife 4, e05178.

Höhener, T.C., Landolt, A., Dessauges, C., Gagliardi, P.A., and Pertz, O. (2022). LITOS - a versatile LED illumination tool for optogenetic stimulation. BioRxiv.

Huang, C.Y., and Ferrell, J.E. (1996). Ultrasensitivity in the mitogen-activated protein kinase cascade. Proc Natl Acad Sci USA 93, 10078–10083.

Hubaud, A., and Pourquié, O. (2014). Signalling dynamics in vertebrate segmentation. Nat. Rev. Mol. Cell Biol. 15, 709–721.

Huebner, R.J., Neumann, N.M., and Ewald, A.J. (2016). Mammary epithelial tubes elongate through MAPK-dependent coordination of cell migration. Development 143, 983–993.

Inman, J.L., Robertson, C., Mott, J.D., and Bissell, M.J. (2015). Mammary gland development: cell fate specification, stem cells and the microenvironment. Development 142, 1028–1042.

Isakoff, S.J., Engelman, J.A., Irie, H.Y., Luo, J., Brachmann, S.M., Pearline, R.V., Cantley, L.C., and Brugge, J.S. (2005). Breast cancer-associated PIK3CA mutations are oncogenic in mammary epithelial cells. Cancer Res. 65, 10992–11000.

Jaqaman, K., Loerke, D., Mettlen, M., Kuwata, H., Grinstein, S., Schmid, S.L., and Danuser, G. (2008). Robust single-particle tracking in live-cell time-lapse sequences. Nat. Methods 5, 695–702.

Kholodenko, B.N., Hancock, J.F., and Kolch, W. (2010). Signalling ballet in space and time. Nat. Rev. Mol. Cell Biol. 11, 414–426.

Kim, N., Kim, J.M., Lee, M., Kim, C.Y., Chang, K.-Y., and Heo, W.D. (2014). Spatiotemporal control of fibroblast growth factor receptor signals by blue light. Chem. Biol. 21, 903–912.

Lam, A.J., St-Pierre, F., Gong, Y., Marshall, J.D., Cranfill, P.J., Baird, M.A., McKeown, M.R., Wiedenmann, J., Davidson, M.W., Schnitzer, M.J., et al. (2012). Improving FRET dynamic range with bright green and red fluorescent proteins. Nat. Methods 9, 1005–1012.

Lauring, J., Cosgrove, D.P., Fontana, S., Gustin, J.P., Konishi, H., Abukhdeir, A.M., Garay, J.P., Mohseni, M., Wang, G.M., Higgins, M.J., et al. (2010). Knock in of the AKT1 E17K mutation in human breast epithelial cells does not recapitulate oncogenic PIK3CA mutations. Oncogene 29, 2337–2345.

Lavoie, H., Gagnon, J., and Therrien, M. (2020). ERK signalling: a master regulator of cell behaviour, life and fate. Nat. Rev. Mol. Cell Biol. 21, 607–632.

Liu, J.S., Farlow, J.T., Paulson, A.K., Labarge, M.A., and Gartner, Z.J. (2012). Programmed cell-to-cell variability in Ras activity triggers emergent behaviors during mammary epithelial morphogenesis. Cell Rep. 2, 1461–1470.

Mailleux, A.A., Overholtzer, M., Schmelzle, T., Bouillet, P., Strasser, A., and Brugge, J.S. (2007). BIM regulates apoptosis during mammary ductal morphogenesis, and its absence reveals alternative cell death mechanisms. Dev. Cell 12, 221–234.

McQuin, C., Goodman, A., Chernyshev, V., Kamentsky, L., Cimini, B.A., Karhohs, K.W., Doan, M., Ding, L., Rafelski, S.M., Thirstrup, D., et al. (2018). CellProfiler 3.0: Next-generation image processing for biology. PLoS Biol. 16, e2005970.

Myers, M.G., Sun, X.J., Cheatham, B., Jachna, B.R., Glasheen, E.M., Backer, J.M., and White, M.F. (1993). IRS-1 is a common element in insulin and insulin-like growth factor-I signaling to the phosphatidylinositol 3’-kinase. Endocrinology 132, 1421–1430.

Paine, I.S., and Lewis, M.T. (2017). The Terminal End Bud: the Little Engine that Could. J. Mammary Gland Biol. Neoplasia 22, 93–108.

Patterson, K.I., Brummer, T., O’Brien, P.M., and Daly, R.J. (2009). Dual-specificity phosphatases: critical regulators with diverse cellular targets. Biochem. J. 418, 475–489.

Pearson, G.W., and Hunter, T. (2007). Real-time imaging reveals that noninvasive mammary epithelial acini can contain motile cells. J. Cell Biol. 179, 1555–1567.

Reginato, M.J., Mills, K.R., Becker, E.B.E., Lynch, D.K., Bonni, A., Muthuswamy, S.K., and Brugge, J.S. (2005). Bim regulation of lumen formation in cultured mammary epithelial acini is targeted by oncogenes. Mol. Cell. Biol. 25, 4591–4601.

Regot, S., Hughey, J.J., Bajar, B.T., Carrasco, S., and Covert, M.W. (2014). High-sensitivity measurements of multiple kinase activities in live single cells. Cell 157, 1724–1734.

Sakaue-Sawano, A., Yo, M., Komatsu, N., Hiratsuka, T., Kogure, T., Hoshida, T., Goshima, N., Matsuda, M., Miyoshi, H., and Miyawaki, A. (2017). Genetically encoded tools for optical dissection of the mammalian cell cycle. Mol. Cell 68, 626–640.e5.

Schafer, Z.T., Grassian, A.R., Song, L., Jiang, Z., Gerhart-Hines, Z., Irie, H.Y., Gao, S., Puigserver, P., and Brugge, J.S. (2009). Antioxidant and oncogene rescue of metabolic defects caused by loss of matrix attachment. Nature 461, 109–113.

Sebastian, J., Richards, R.G., Walker, M.P., Wiesen, J.F., Werb, Z., Derynck, R., Hom, Y.K., Cunha, G.R., and DiAugustine, R.P. (1998). Activation and function of the epidermal growth factor receptor and erbB-2 during mammary gland morphogenesis. Cell Growth Differ. 9, 777–785.

Shaulian, E., and Karin, M. (2001). AP-1 in cell proliferation and survival. Oncogene 20, 2390–2400.

Shcherbakova, D.M., Baloban, M., Emelyanov, A.V., Brenowitz, M., Guo, P., and Verkhusha, V.V. (2016). Bright monomeric near-infrared fluorescent proteins as tags and biosensors for multiscale imaging. Nat. Commun. 7, 12405.

Sparta, B., Pargett, M., Minguet, M., Distor, K., Bell, G., and Albeck, J.G. (2015). Receptor Level Mechanisms Are Required for Epidermal Growth Factor (EGF)-stimulated Extracellular Signal-regulated Kinase (ERK) Activity Pulses. J. Biol. Chem. 290, 24784–24792.

Sternlicht, M.D., Sunnarborg, S.W., Kouros-Mehr, H., Yu, Y., Lee, D.C., and Werb, Z. (2005). Mammary ductal morphogenesis requires paracrine activation of stromal EGFR via ADAM17-dependent shedding of epithelial amphiregulin. Development 132, 3923–3933.

Tikoo, A., Roh, V., Montgomery, K.G., Ivetac, I., Waring, P., Pelzer, R., Hare, L., Shackleton, M., Humbert, P., and Phillips, W.A. (2012). Physiological levels of Pik3ca(H1047R) mutation in the mouse mammary gland results in ductal hyperplasia and formation of ERα-positive tumors. PLoS ONE 7, e36924.

Valon, L., Davidović, A., Levillayer, F., Villars, A., Chouly, M., Cerqueira-Campos, F., and Levayer, R. (2021). Robustness of epithelial sealing is an emerging property of local ERK feedback driven by cell elimination. Dev. Cell 56, 1700–1711.e8.

Wait, E., Winter, M., Bjornsson, C., Kokovay, E., Wang, Y., Goderie, S., Temple, S., and Cohen, A.R. (2014). Visualization and correction of automated segmentation, tracking and lineaging from 5-D stem cell image sequences. BMC Bioinformatics 15, 328.

Wait, E., Winter, M., and Cohen, A.R. (2019). Hydra image processor: 5-D GPU image analysis library with MATLAB and python wrappers. Bioinformatics 35, 5393–5395.

Wang, H., Lacoche, S., Huang, L., Xue, B., and Muthuswamy, S.K. (2013). Rotational motion during three-dimensional morphogenesis of mammary epithelial acini relates to laminin matrix assembly. Proc Natl Acad Sci USA 110, 163–168.

Winter, M.R., Fang, C., Banker, G., Roysam, B., and Cohen, A.R. (2012). Axonal transport analysis using Multitemporal Association Tracking. Int. J. Comput. Biol. Drug Des. 5, 35–48.

Winter, M., Wait, E., Roysam, B., Goderie, S.K., Ali, R.A.N., Kokovay, E., Temple, S., and Cohen, A.R. (2011). Vertebrate neural stem cell segmentation, tracking and lineaging with validation and editing. Nat. Protoc. 6, 1942–1952.

Winter, M., Mankowski, W., Wait, E., Temple, S., and Cohen, A.R. (2016). LEVER: software tools for segmentation, tracking and lineaging of proliferating cells. Bioinformatics 32, 3530–3531.

Winter, M., Mankowski, W., Wait, E., De La Hoz, E.C., Aguinaldo, A., and Cohen, A.R. (2019). Separating touching cells using pixel replicated elliptical shape models. IEEE Trans. Med. Imaging 38, 883–893.

Young, C.D., Zimmerman, L.J., Hoshino, D., Formisano, L., Hanker, A.B., Gatza, M.L., Morrison, M.M., Moore, P.D., Whitwell, C.A., Dave, B., et al. (2015). Activating PIK3CA Mutations Induce an Epidermal Growth Factor Receptor (EGFR)/Extracellular Signal-regulated Kinase (ERK) Paracrine Signaling Axis in Basal-like Breast Cancer. Mol. Cell. Proteomics 14, 1959–1976.

Yusa, K., Zhou, L., Li, M.A., Bradley, A., and Craig, N.L. (2011). A hyperactive piggyBac transposase for mammalian applications. Proc Natl Acad Sci USA 108, 1531–1536.

