## Supplemental Material for "Spatio-temporal Control of ERK Pulse Frequency Coordinates Fate Decisions during Mammary Acinar Morphogenesis"

**Supplemental Text**

We used our optogenetic system to explore different hypothetical mechanisms of the spatio-temporal regulation of ERK pulse frequency and its interpretation into fate decisions. We first explored if the MAPK network reacts differently to extracellular inputs in outer and inner cells in stage 2 acini. The MAPK pathway is wired to produce ultrasensitive, all-or-nothing ERK pulses at threshold GF inputs (Huang and Ferrell 1996). In between the input range that does not activate ERK, and the one that produces robust full amplitude ERK pulses, a small input range often leads to non-robust, heterogeneous ERK responses (Ryu et al. 2015). We therefore reasoned that differential ERK pulse frequency observed in outer/inner cells might result from the ability of the MAPK network to shift its sensitivity to inputs depending on its location within an acinus. We therefore identified optoFGFR light inputs of different strengths that either do not activate ERK, that lead to ERK pulses with heterogeneous amplitudes, or lead to full amplitude ERK pulses (Figure S5A). We observed that the different light-induced optoFGFR inputs of different strengths, which are perceived equally across all the cells of the acinus, led to identical ERK responses irrespective of the location of single cells within stage 2 acini. This suggests that the ERK pulse frequency does not depend on spatial regulation of the MAPK network's ability to produce switch-like, all-or-nothing ERK responses.

Second, we reasoned that a 1st receptor input might desensitize the MAPK network to a 2nd input differently in inner and outer cells in stage 2 acini, as a possible mechanism of spatio-temporal control of ERK pulse frequency. This could for example be regulated by ERK-dependent transcriptional activation of dual specificity phosphatases (DUSPs) that switch off ERK signaling. To test this hypothesis, we stimulated stage 2 acini with successive optoFGFR inputs delivered at 15- and 21-minutes intervals. These correspond to specific times at which an ERK pulse has not yet adapted to baseline (15 minutes interval), and at which an ERK pulse has adapted to the baseline. We observed that these specific temporal stimulation schemes did not yield different ERK dynamics in inner and outer cells in stage 2 acini (Figure S5B). This strongly suggests the absence of spatio-temporal control of ERK-dependent desensitization of ERK pulses in stage 2 acini.

### References

Huang, C.Y., and Ferrell, J.E. (1996). Ultrasensitivity in the mitogen-activated protein kinase cascade. Proc Natl Acad Sci USA 93, 10078–10083.

Ryu, H., Chung, M., Dobrzyński, M., Fey, D., Blum, Y., Lee, S.S., Peter, M., Kholodenko, B.N., Jeon, N.L., and Pertz, O. (2015). Frequency modulation of ERK activation dynamics rewires cell fate. Mol. Syst. Biol. 11, 838.

### Supplemental Figures

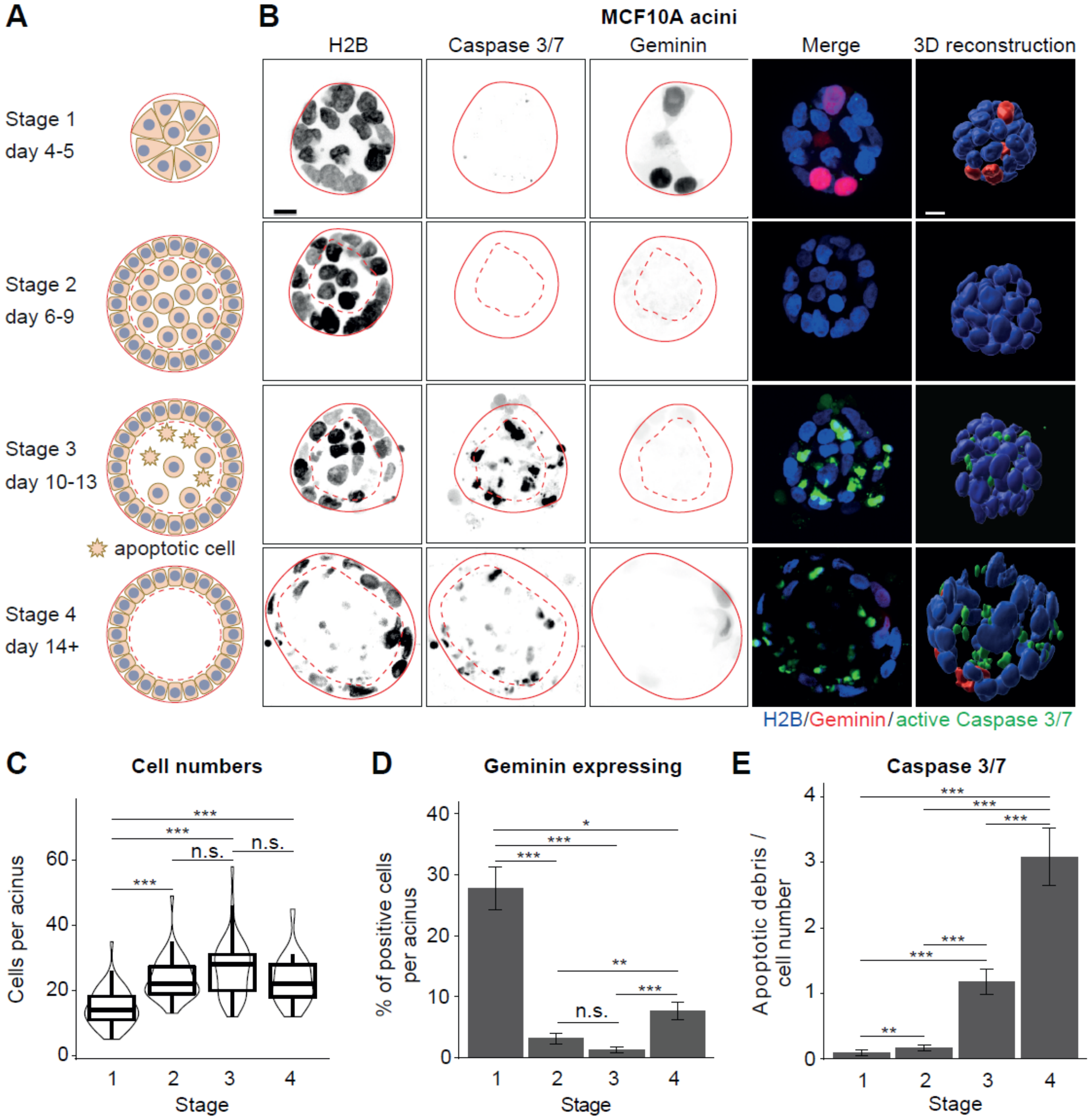


Figure S1 **Distinct stages of MCF10A acinar morphogenesis.** (A) Schematics of acinar morphogenetic stages. (B) Micrographs and 3D reconstructions of H2B, caspase 3/7 fluorogenic substrate and geminin signals in acini corresponding to the stages in (A). Micrographs show maximal intensity projections of equatorial Z planes spanning 12 µm. Plain lines mark the borders of the acini, dashed lines mark the outer cell layer. Scale bar = 10 µm. (C) Cell numbers per acinus at the different stages (N = 28 - 60 acini each). (D) Fraction of Geminin positive cells per acinus at different stages. Same acini as in (C). (E) Number of Caspase 3/7 apoptotic debris divided by the acinar cell number at different stages. Same acini as in (C). (D-E) Error bars represent standard error of the mean. (C-E) Wilcoxon tests (n.s., $P > 0.05$; *, $P < 0.05$; **, $P < 0.01$; ***, $P < 0.001$).

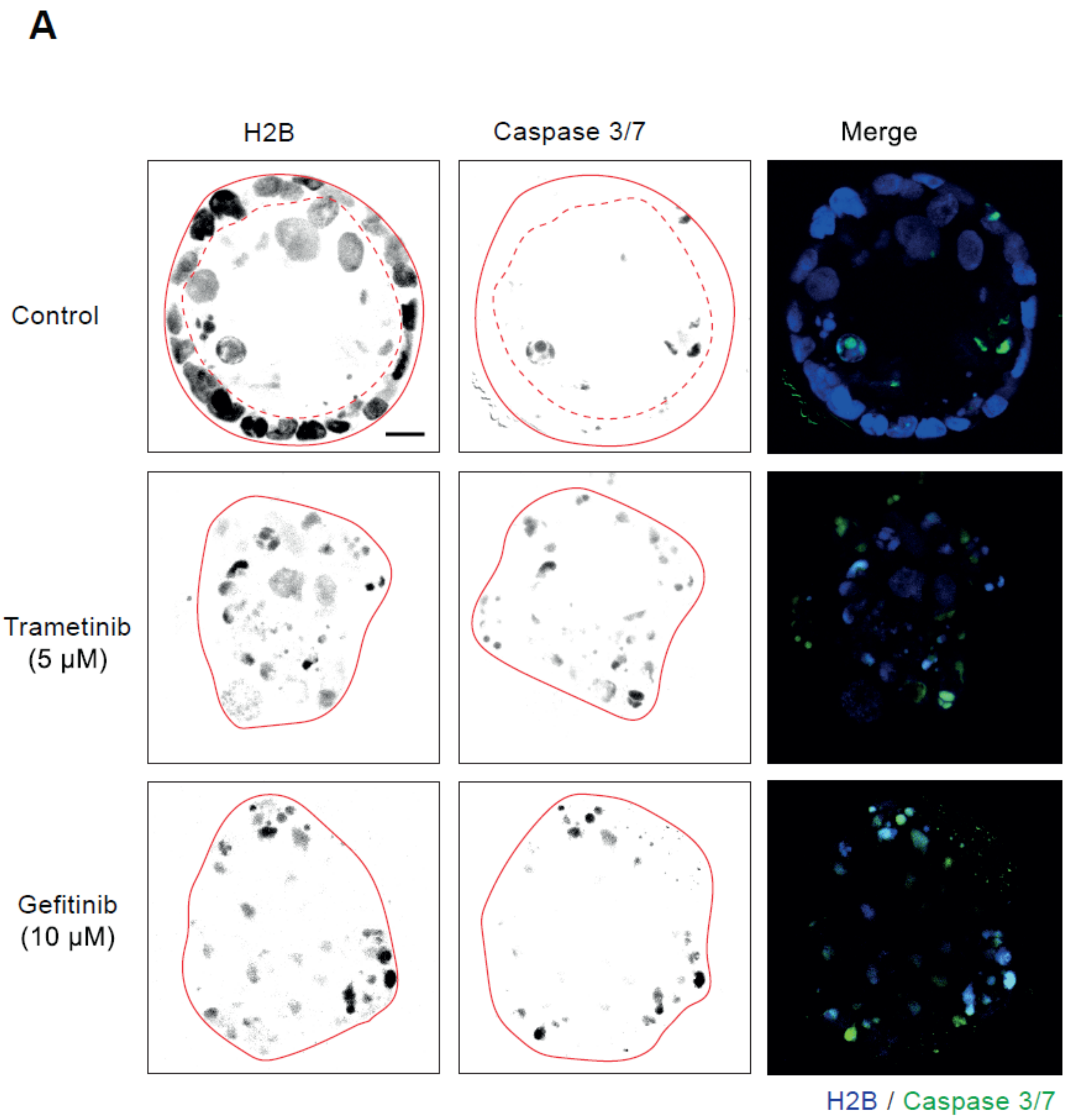


Figure S2 **Acini culture conditions and effect of blocking ERK pulses on cell survival.** Micrographs of H2B and caspase 3/7 fluorogenic substrate in stage 4 control acini and acini treated with Trametinib or Gefitinib. Micrographs show maximal intensity projections of equatorial Z planes spanning 12 µm. Plain lines mark the borders of the acini, dashed lines mark the outer cell layer. Scale bar = 10 µm.

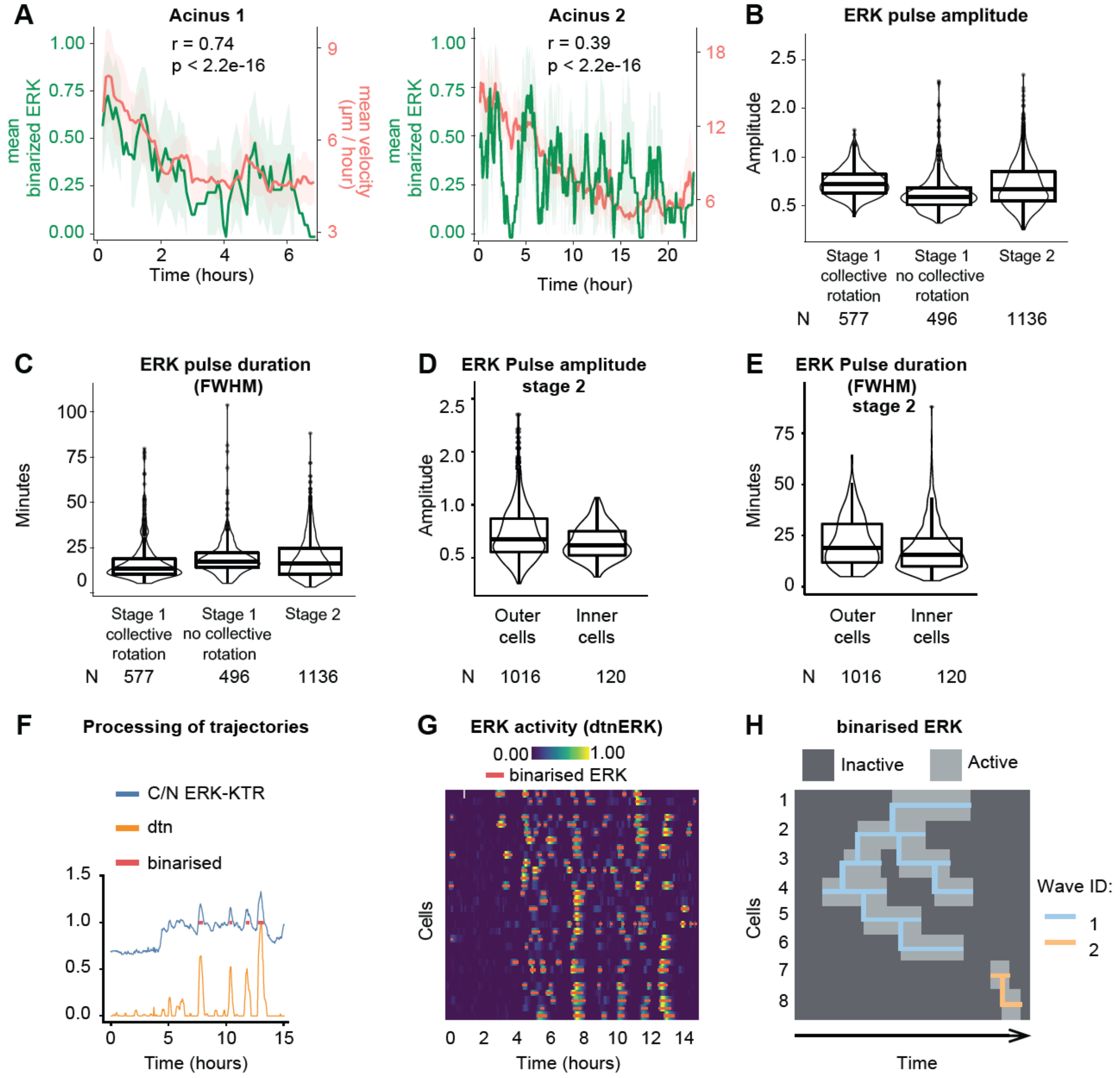


Figure S3 **Additional experiments and quantification of ERK dynamics properties during acinar morphogenesis.** (A) Analysis of motility and ERK activity in 2 different acini. Graphs show mean binarized ERK activity and mean instantaneous velocity with 95% confidence intervals of all imaged cells over time and their Pearson correlation coefficient. Mean binarized ERK activity is used as a measure for the fraction of the cell population in a state of active ERK. (B) ERK pulse amplitudes from trajectories at different developmental timepoints. Same tracks as analyzed in Fig. 2D. (C) ERK pulse durations from trajectories at different developmental timepoints. Same tracks as analyzed in Fig. 2D. (D) ERK pulse amplitudes from trajectories of cells located on the outer versus the inner layers of stage 2 acini. Same tracks as analyzed in Fig. 3A. (E) ERK pulse durations from trajectories of cells located on the outer versus the inner layers of stage 2 acini. Same tracks as analyzed in Fig. 3A. (F) Raw, detrended and normalized (dtn) and binarized ERK activity trajectories in a cell from a stage 2 acini. Binarized ERK activity was used for detection of collective events. (G) Heatmap color-coded dtn ERK activity time series in single cells, overlayed with the binarized signal (red = ERK on). (H) Identification and tracking of collective events based on the binarized ERK activity signal. First occurrences of ERK activity are detected and linked to subsequent ERK activity in neighboring cells.

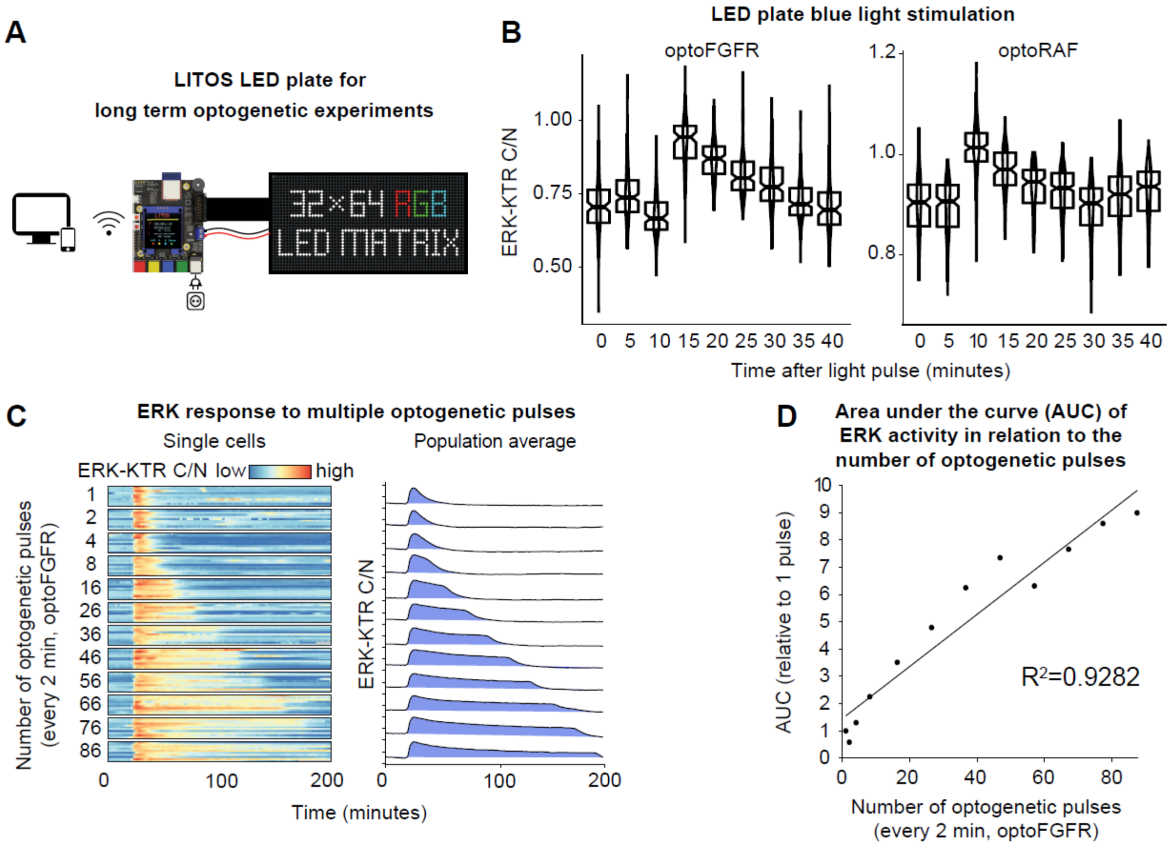


Figure S4 **Optogenetic stimulation of acini with a programmable LED plate and calibration of area under the curve (AUC) vs number of pulses**. (A) Scheme of the LITOS system used for long-term optogenetic stimulation of acini. (B) ERK activity levels in cells from optoFGFR and optoRAF acini that were fixed at the indicated time points after a 1 minute stimulation with blue light from the LED plate system. (C) MCF10A cells stably expressing the optogenetic actuator optoFGFR and the ERK-KTR-mRuby2 biosensor were stimulated with repeated blue light pulses to induce longer ERK pulses with larger AUC. Both blue light pulse frequency and image acquisition were set at 2 minute frequency. Different amounts of repeated blue light pulses were used to induce different AUC. Randomly selected single-cell trajectories are represented in color code on the left. Average ERK activity on all the cells in the captured field of view with AUC in blue is shown on the right. (D) Relation of AUC in respect to the number of consecutive blue light pulses modeled by linear regression.

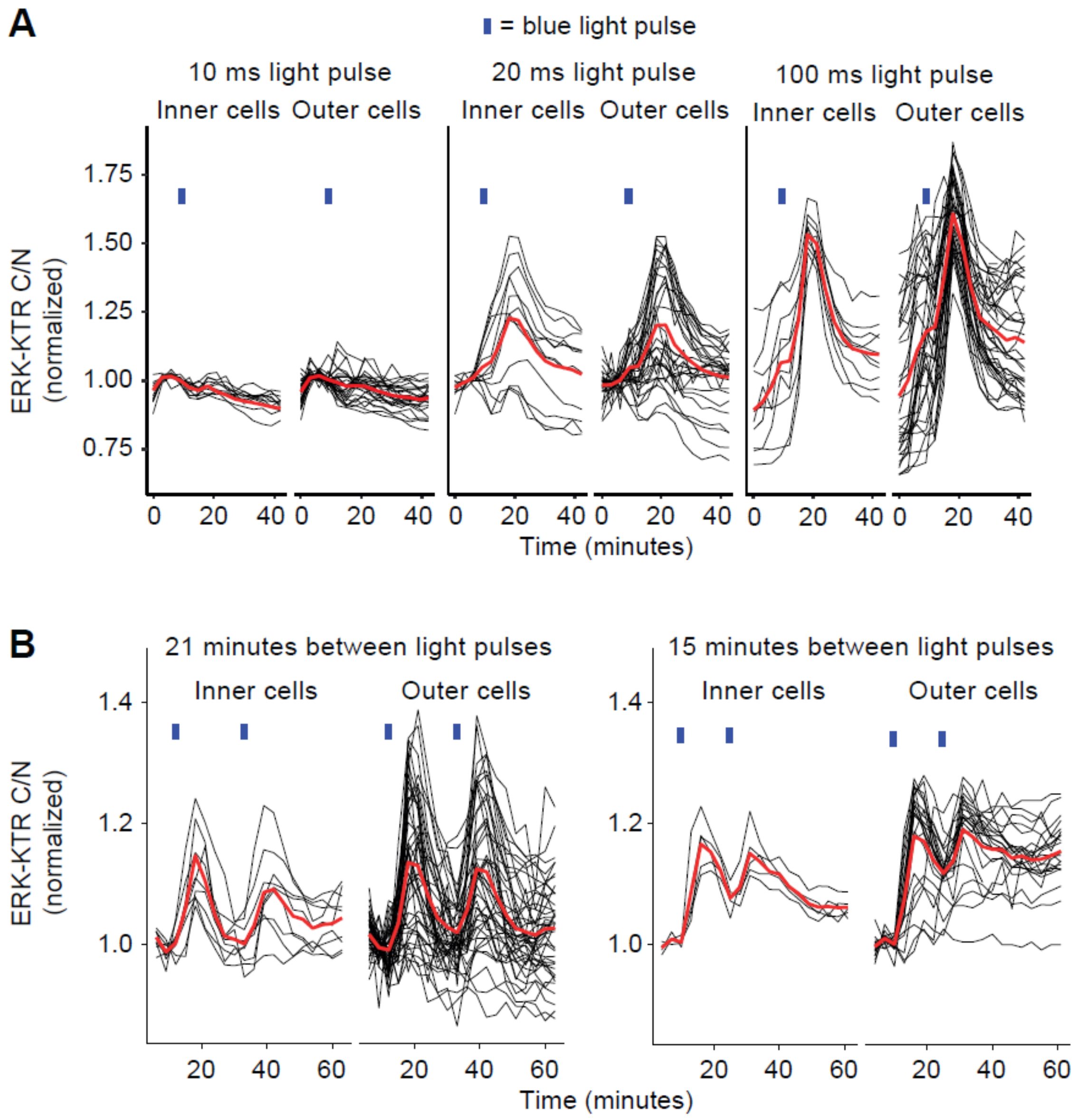


Figure S5 **ERK pulse threshold and refractory period in inner and outer acinar cells.** (A) Single cell ERK activity trajectories from an optoFGFR expressing acinus stimulated with a blue light pulse of varying exposure time after 10 minutes of imaging under the microscope. The same acinus was used for all experiments. (B) Single cell ERK activity trajectories from optoFGFR expressing acini stimulated with 2 subsequent blue light pulses with a 15 or 21 minute interval between the pulses.

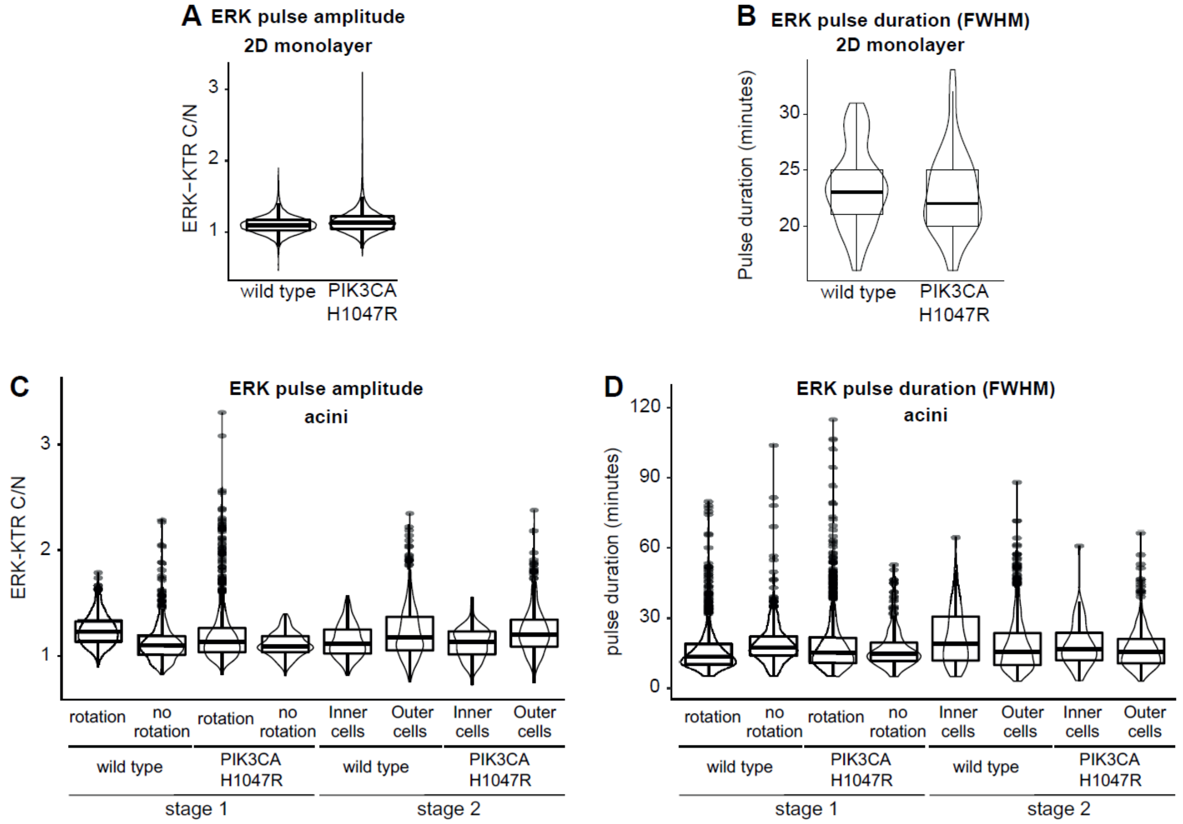


Figure S6 **Quantification of ERK pulse amplitude and width in WT and PIK3CA H1047R MCF10A monolayer cells and acini.** (A) ERK pulse amplitudes in WT and PIK3CA H1047R monolayer cells. (B) ERK pulse durations in WT and PIK3CA H1047R monolayer cells. (C) ERK pulse amplitudes from trajectories of WT and PIK3CA H1047R cells at different stages and locations within the acinus. Same tracks as analyzed in Fig. 7B. (D) ERK pulse durations from trajectories of WT and PIK3CA H1047R cells at different stages and locations within the acinus. Same tracks as analyzed in Fig. 7B.

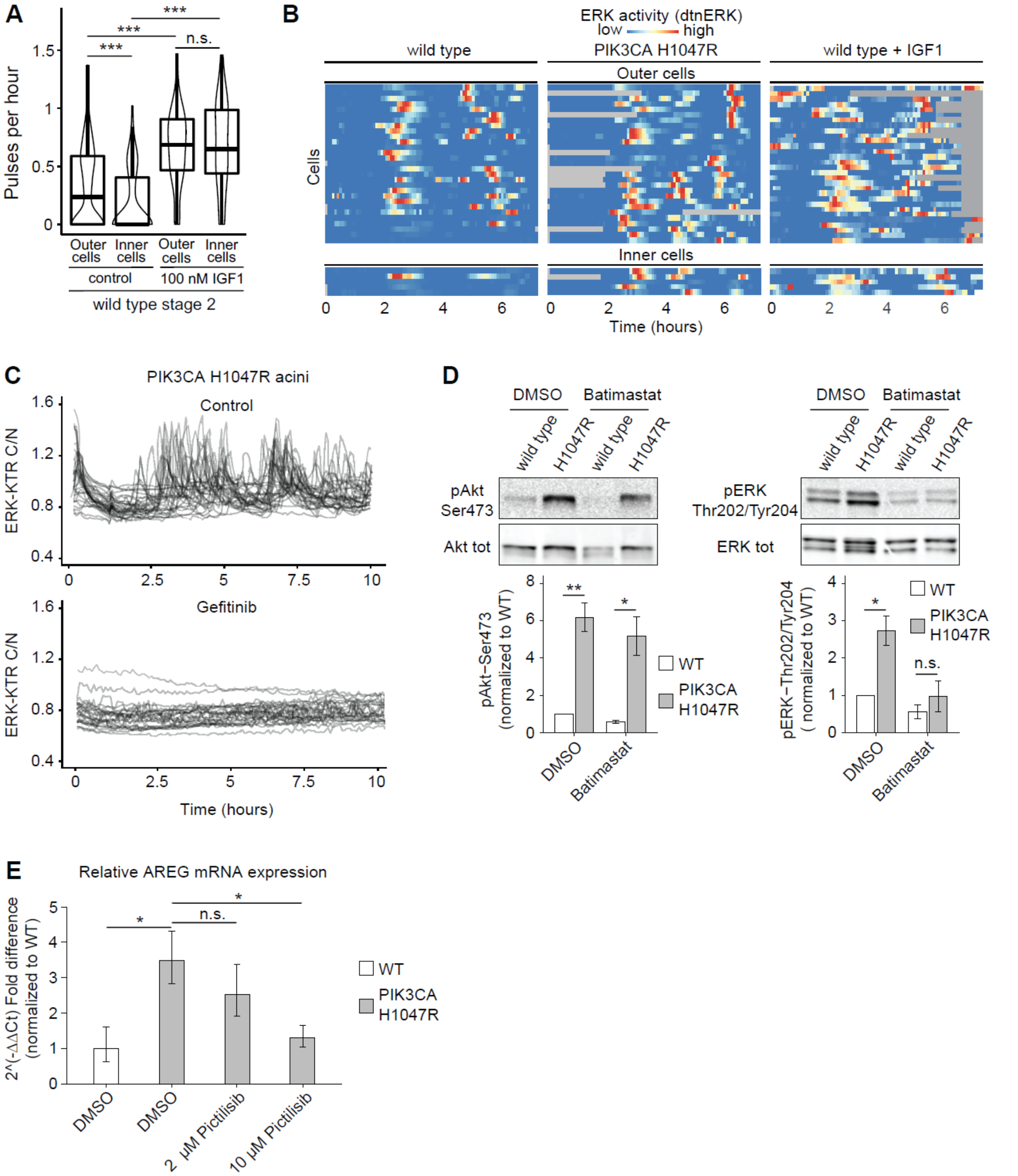


Figure S7 **PIK3CA H1047R induces increase of ERK pulse frequency via a MMP-EGFR mediated mechanism.** (A) ERK frequencies from control acini and acini treated with 100 nM IGF1. Control data is the same as in Figure 3A. Wilcoxon tests (n.s., P > 0.05; ***, P < 0.001). (B) Heatmaps of detrended and normalized single cell ERK activity levels over time in outer and inner cells of representative WT stage 2 acini, PIK3CA H1047R stage 2 acini and IGF1-treated WT stage 2 acini. (C) Overlayed single cell ERK activity trajectories from control and gefitinib treated PIK3CA H1047R acini. (D) Representative blots of phospho-AKT$^{Ser473}$ (pAKT $^{Ser473}$), p44/42 MAPK

(Erk1/2)$^{\text{Thr202/Tyr204}}$, AKT, p44/42 MAPK (Erk1/2)$^{\text{Thr202/Tyr204}}$, and p44/42 MAPK (Erk1/2) are presented. Quantification of protein abundances of vehicle (DMSO) or Batimastat treated WT and PIK3CA H1047R MCF10A cells is shown relative to the levels in vehicle treated WT cells. Data from 4 independent experiments. Paired t-test (n.s., $P > 0.05$; *, $P < 0.05$; **, $P < 0.01$). (E) The expression of EGFR ligand AREG in WT and PIK3CA H1047R mutant MCF10A cells cultured in starvation media and treated with vehicle (DMSO) or indicated concentration of pictilisib was determined by qPCR and normalized to the expression of FLJ22101; the normalized expression of AREG is presented relative to WT group (average of technical triplicates ± STD) Paired t-test (n.s., $P > 0.05$; *).

**Supplemental Movies**

Movie S1 **Transition from collective rotational motility to low motility state in a stage 1 acinus.** 3D reconstruction of a stage 1 MCF10A acinus cross section. Tracks from 3 cells (indicated by gray spheres) are shown. Tracks are color-coded for instantaneous velocity.

Movie S2 **Transition from high to low ERK frequency in a stage 1 acinus.** LEVERJS 3D reconstruction of a stage 1 MCF10A acinus cross section. The cells with track IDs 4 and 7 are labeled as examples. Top left panel: H2B - miRFP channel with nuclear segmentation overlaid as wireframes. Wireframes are colored by track ID. Bottom left panel: Nuclear segmentations are color-coded by detrended and normalized ERK activity. Dark/bright gray shades indicate low/high ERK activities. Middle panels: ERK-KTR–mTurquoise2 channel. Optical sections are focused on the centroids of cells 4 (top) and 7 (bottom). Right panels: detrended and normalized ERK trajectories of cells 4 (top) and 7 (bottom) with the time axis labeled in frames.

Movie S3 **Collective ERK waves in a stage 2 acinus.** LEVERJS 3D reconstruction of a stage 2 MCF10A acinus. The cell with track ID 6 is labeled as an example. Top left: H2B-miRFP channel. Bottom left: H2B- miRFP channel with nuclear segmentation overlaid as wireframes. Wireframes are colored by track IDs. Top middle: ERK-KTR – mTurquoise2 channel. Optical sections are focused on cell 6. Bottom middle: Nuclear segmentations are color-coded by detrended and normalized ERK activity. Dark/bright gray shades indicate low/high ERK activities. Right: detrended and normalized ERK activity trajectory of cell 6 with the time axis labeled in frames.

Movie S4 **Optogenetic entrainment of ERK frequency in stage 1 rotating acini.** Representative day 4 acinus expressing H2B – miRFP (left), ERK-KTR (right) and optoFGFR. The acinus was imaged every 5 minutes for a total time of 8 hours. In the initial 4 hours no optogenetic stimulation was applied. In the subsequent 4 hours, the acinus was stimulated with blue light every 30 minutes. Nuclei were segmented with Imaris and migration tracks were color-coded according to instantaneous speed (left). Time points of blue light stimulation are indicated by a cyan bar above the images. Scale bar: 10 µm.

Movie S5 **Optogenetic entrainment of ERK frequency in stage 2 acini.** LEVERJS 3D reconstructions of stage 2 MCF10A acini expressing optoFGFR that were stimulated with blue light pulses at different frequencies. Top: H2B - miRFP channel with nuclear segmentation overlaid as wireframes. Wireframes are colored by track ID. Bottom: Nuclear segmentations are color-coded by detrended and normalized ERK activity. Dark/bright gray shades indicate low/high ERK activities. Time points of blue light stimulation are indicated by a blue signal in the bottom panel.

Movie S6 **ERK pulses in stage 2 WT versus PIK3CA H1047R acini.** LEVERJS 3D reconstructions of stage 2 wild type and PIK3CA H1047R MCF10A acinus cross sections side by side. Top: H2B-miRFP channel with nuclear segmentation overlaid as wireframes. Wireframes are colored by track ID. Bottom: Nuclear segmentations are color-coded by detrended and normalized ERK activity. Dark/bright gray shades indicate low/high ERK activities.
